# Reinforcement-based option competition in human dorsal stream during exploration/exploitation of a continuous space

**DOI:** 10.1101/2023.05.22.541828

**Authors:** Michael N. Hallquist, Kai Hwang, Beatriz Luna, Alexandre Y. Dombrovski

**Affiliations:** Department of Psychology, University of North Carolina, Chapel Hill, NC, USA; Department of Psychological and Brain Sciences, Iowa Neuroscience Institute, University of Iowa, Iowa City, IA, USA; Department of Psychiatry, University of Pittsburgh, Pittsburgh, PA, USA

## Abstract

Primates exploring and exploiting a continuous sensorimotor space rely on maps in the dorsal stream that guide visual search, locomotion, and grasp. For example, an animal swinging from one tree limb to the next uses rapidly evolving sensorimotor representations to decide when to harvest a reward. We show that such exploration/exploitation depends on dynamic maps of competing option values in the human dorsal stream. Using a reinforcement learning (RL) model capable of rapid learning and efficient exploration and exploitation, we show that preferred options are selectively maintained on the map while the values of spatiotemporally distant alternatives are compressed. Consistent with biophysical models of cortical option competition, dorsal stream BOLD signal increased and posterior cortical β_1_/α oscillations desynchronized as the number of potentially valuable options grew, matching predictions of information-compressing RL rather than traditional RL that caches long-term values. BOLD and β_1_/α responses were correlated and predicted the successful transition from exploration to exploitation. These option competition dynamics were observed across parietal and frontal dorsal stream regions, but not in the occipito-temporal MT+ sensitive to the average reward rate. Our results also illustrate that models’ diverging predictions about information dynamics can help to adjudicate between them based on population activity.

**Graphical abstract:** 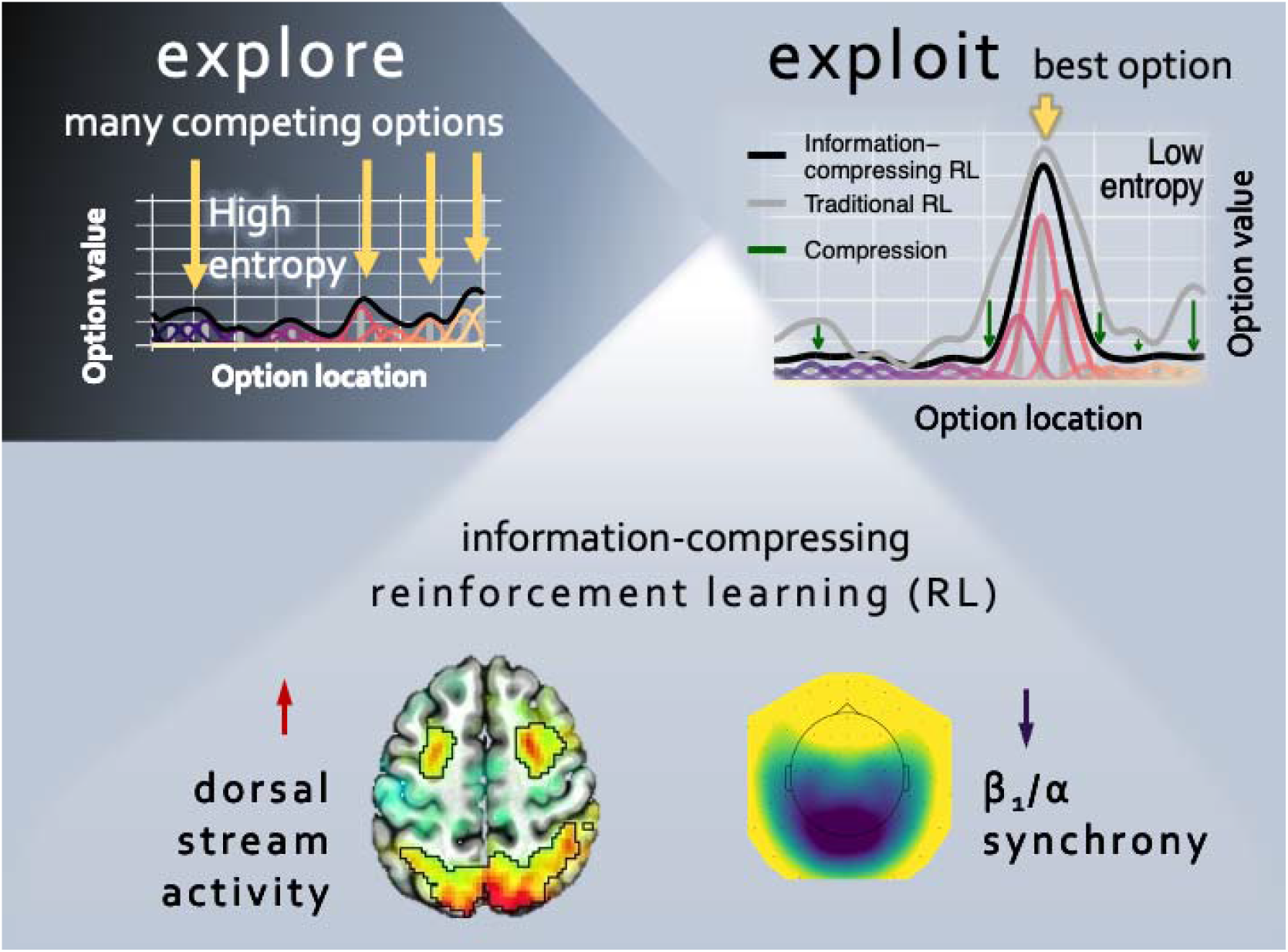

## Introduction

Organisms face a difficult dilemma between exploiting options that are known to be good and exploring new options that might be even better. When a vertebrate faces a few discrete options, the striatum and amygdala can resolve the explore-exploit dilemma by representing options egocentrically (e.g., right/left) and tuning the exploration rate based on meso-striatal dopaminergic signals ^1–4^. In more complex terrestrial environments, however, quadrupeds rely on world-centric hippocampal cognitive maps that incorporate reinforcement and are stored in long-term memory ^5–7^. While these mechanisms work efficiently at slower timescales, exploration and exploitation become more demanding when we move rapidly through a dynamic environment.

For example, as an adaptation to arboreal hunting and foraging on terminal branches, primates evolved visuo-motor systems that support fast and precise visually guided actions ^8^. These behaviors rely on the cortical “where” stream, or the dorsal attention network (DAN), which integrates visual and somatosensory information to build dynamic world-centric maps that guide visual search, locomotion, and grasp. More specifically, the posterior parietal cortex (PPC) constructs maps using visual inputs from temporal-occipital areas such as MT+ as well as parietal somatosensory inputs. In turn, the PPC sends map-based outputs to the frontal dorsal and ventral premotor (PMd and PMv) cortex and the frontal eye fields (FEF) that guide action ^9–11^.

Visuomotor learning has to occur at a faster timescale than instrumental learning in the basal forebrain and striatum, which integrates reinforcement slowly and retains long-term values ^12^. Moreover, PPC maps contain rich visuomotor data necessary to decide what actions are likely to succeed in current and upcoming spatio-temporal locations, e.g., when an insect can be grasped on a moving branch ^13–15^. These pragmatic maps represent programs of movement toward currently available options that are based on prior visuomotor experience^9^. Studies of gaze control, for example, find that PPC facilitates goal-congruent saccades by comparing what one is looking at versus what one is looking for^10^.

How are these goals set? The PPC integrates past visuomotor experience and rewards ^16^, establishing bi-directional links between attention and learning ^15^. Visual stimuli repeatedly paired with rewards gradually gain priority on the PPC map and will be preferred in visuomotor interactions such as grasping actions. When a primate faces an array of potentially valuable options, PPC subpopulations representing them compete for behavioral selection^9^. Yet, we do not understand the mechanisms that enable the rapid integration of reinforcement into PPC maps. This is in part because most studies of reward learning employed a handful of spatially unstructured options that may require little involvement of the PPC.

Here, we considered how encoding of reinforcement in the PPC, occipito-temporal and prefrontal DAN regions may resolve option competition and enable exploitation. We experimentally manipulated the distribution of rewards during rapid movement through a one-dimensional continuous space, as a clock hand revolves around a circle (Figure 1A), inducing value-laden continuous visuomotor representations. We tested the general hypothesis that representations of reinforcement history in PPC are integrated into a map that supports exploitation of the most-rewarded option. Given the rapid dynamics of visuomotor learning in PPC, prior studies have considered working memory (WM)^17, 18^ or serial hypothesis testing^14, 19^. Leveraging a previously validated computational model ^20^, we demonstrate, however, that option competition in the DAN cannot be fully explained by WM or traditional RL, but involves an information-compressing RL process that selectively maintains the values of preferred options and allows non-preferred alternatives to decay.

**Figure 1.**
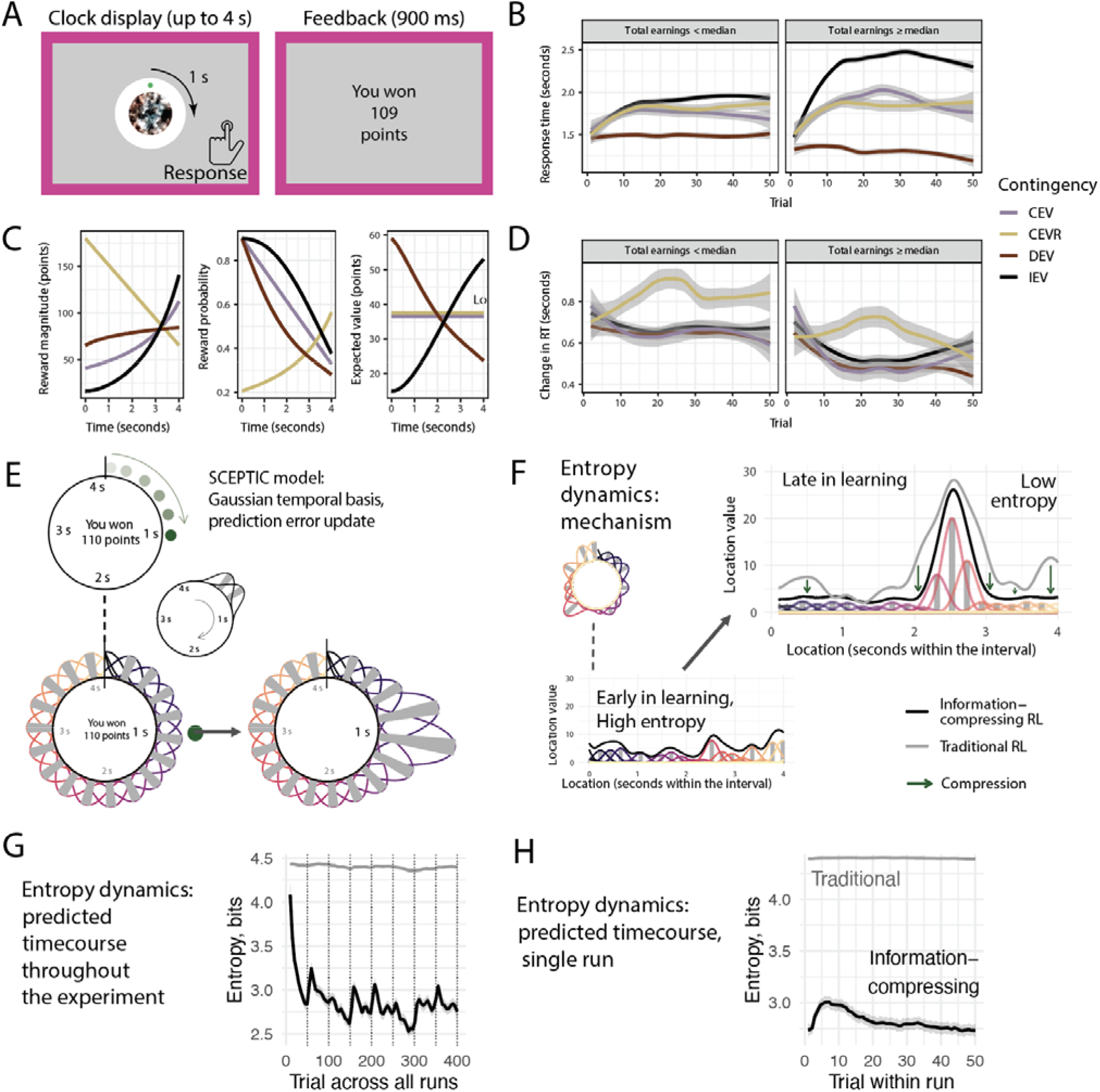
Paradigm and SCEPTIC model. (A) The clock paradigm consists of decision and feedback phases. During the decision phase, a dot revolves 360° arou nd a central stimulus over the course of four seconds. Participants press a button to stop the revolution and receive a probabilistic outcome. (B) Evolution of subjects’ response times (RT) by contingency and performance. Panels represent participants whose total earnings were above or below the sample median. (C) Rewards are drawn from one of four monotonically time-varying contingencies: two learnable (increasing expected value, IEV, and decreasing expected value, DEV, dark colors), and two unlearnable (constant expected value, CEV, and constant expected value–reversed, CEVR, light colors). Reward probabilities and magnitudes vary independently. (D) Evolution of subjects’ response time swings (RT swings) by contingency and performance. (E) SCEPTIC model: basis function representation. Top: Subject responds at 1s and wins 110 points. Bottom left: The 1D space of the task is tiled with Gaussian-shaped learning elements with staggered receptive fields. Bottom right: the reward at 1s updates expected values (weights) of nearby basis elements. (F) Entropy dynamics of the information-compressing model, mechanism. Left: example value distribution early in learning, when all locations have similar values and entropy is high (top: visual space of the task, bottom: linear coordinates with location on abscissa. Right: example value distribution late in learning, contrasting information-compressing RL (black line, colored bases) with traditional RL (grey line). Information compression (green arrows) reduces the entropy of the value distribution. (G) Entropy dynamics of the information-compressing vs. traditional RL model across the entire experiment. High initial entropy reflects random uniform prior basis element weights, however qualitatively the same dynamics are seen with priors of 0 (Figure s1D-F). (H) The same as G, for an average run. First run is excluded to eliminate effect of priors.

The key insight here is that the entropy of values within a map tunes the explore-exploit balance^20^ (cf. policy entropy in artificial intelligence^21^). Consider a learning agent who tracks the expected reward or value associated with each target or option, termed the value function. When the estimated values of all options are equal, the entropy of the value function is highest, and the agent can needs to explore to discovery truly superior options. Conversely, when a single superior option (global value maximum) can be exploited, entropy is low. Thus, entropy of the value function quantifies global uncertainty about which option is best. Exploration generally increases the mutual information between the learned (agent’s) and objective (environment’s) value functions. However, a reward-maximizing agent only needs to discover the highest-valued options, rather than attempting to learn precisely the value of every option^22^. Furthermore, maintaining and updating a detailed map incurs a high memory cost and a risk of cognitive failure, which humans strive to minimize^23^. Thus, a resource-rational agent should reduce the entropy of its learned value function^cf. 24^.

We have previously shown that selectively maintaining the values of preferred options and forgetting non-preferred alternatives reduces the entropy or compresses the information contained in the value function^20^. This information-compressing RL model learns and forgets faster than traditional RL, almost as quickly as classical buffer WM models. Yet, while working memory excels in recognizing deterministic rules, its limited buffer becomes a hindrance in stochastic environments. By contrast, resource-rational information-compressing RL integrates reinforcement over a period sufficiently long to explore and exploit stochastic environments efficiently. We have previously shown in a continuous space that it outperforms more memory-intensive traditional RL with long-term value persistence ^20^. Moreover, whereas WM models fail to explain empirical findings of learned long-term value or salience signals in the PPC ^25, 26^, information-compressing RL accounts for them and makes a key neural prediction: Increases in the entropy of the learned value function, and consequently the number of potentially good options, should recruit more PPC neuronal subpopulations representing them. Entropy decreases should have the opposite effect, as fewer subpopulations dominate the output and behavior shifts from exploration to exploitation. Critically, traditional RL does not predict entropy decreases during successful learning and does not link entropy dynamics to exploitation ^20^. By contrast, WM models predict divergent entropy dynamics, determined only by the content of the buffer.

Here, we adjudicate among alternative accounts of value-based option competition in the DAN: traditional RL, information-compressing RL, and WM, alone or in combination. Biophysical models and electrophysiological studies of option competition in the posterior cortex suggest that each competing option may be encoded by a subpopulation with a unique phase of β_1_/α oscillatory output^17, 18, 27–30^. Thus, we hypothesized that increases in the number of close-valued options would induce a β_1_/α desynchronization. We also examined whether BOLD and oscillatory dynamics consistent with information-compressing RL would predict a successful explore-exploit transition. Two studies of DAN BOLD and one MEG study of posterior oscillations provided evidence supporting information-compressing RL.

## Results

We begin by describing (i) behavior on the clock task and our information-compressing RL model, SCEPTIC (StrategiC Exploration/exploiTation of Instrumental Contingencies) and (ii) the connectivity-based parcellation of the human DAN used here. Our main analyses focus on entropy dynamics and the transition from exploration to exploitation. We report on distinct neural substrates of exploration elsewhere.

### Behavior and SCEPTIC model

On the clock task (Figure 1A), participants explore and learn reward contingencies within a four-second time interval, presenting a challenging unidimensional continuous environment. The passage of time is marked by the rotation of a dot around a clock face, reducing demands on internal timing. They are told to find the response time that yields the most points. To encourage extensive exploration and trial-by-trial learning, the task employs four stochastic reward contingencies with varying reward probability/magnitude tradeoffs (Figure 1C), which require integration of reinforcement over time and impede purely WM-based or heuristic strategies. Indeed, whereas people’s responses shifted toward value maxima in learnable contingencies (Figure 1B), even more successful participants tended not to respond as early as possible in DEV or as late as possible in IEV. Thus, participants generally did not recognize that contingencies were monotonic, instead searching for a subjective value maximum (RT _Vmax_); their estimate of its location often shifted within the block. Trial-wise changes of response times (aka ‘RT swings’) provide a model-free index of exploration. Early in learning, better-performing participants displayed large RT swings followed by a decline as they shifted to exploiting the subjective value maximum. Less successful participants kept exploring stochastically, with moderately large RT swings throughout, never settling on a clear value maximum. Curiously, successful participants transitioned from exploration to exploitation even in unlearnable contingencies where no objective value maximum exists ^20^. As detailed in the next section, this behavior can be explained by adaptive selective maintenance of reinforcement histories.

Our SCEPTIC reinforcement learning model ^20^ quantifies both local reinforcement (reward prediction errors) and global value map updates. On the clock task SCEPTIC approximates the value function or expected reward across the space (interval) with a set of learning elements whose temporal receptive fields cover the interval^31, 32^. Each element learns from temporally proximal rewards, updating its weight by reward prediction errors or the discrepancy between model-predicted reward at the chosen RT and the obtained reward (Figure 1E). The highest-valued RT is the global maximum (aka RT _Vmax_) of the model-estimated value function (Figure 1F).

To understand how global map updates can be quantified, we can think of locations (basis elements) as an alphabet, reinforcement sequences as messages, and the value function as an information source encoding the reinforcement history. The information content of this source is given by Shannon’s entropy of the value function (normalized element weights), which is high when multiple attractive options compete and low when a single option dominates (Figure 1F). Thus, increases in entropy reflect the emergence of competing options on the global map. We have previously found that human behavior on the clock task is best explained by a model that selectively maintains the values of favored actions and allows the alternatives to decay, compressing the information content – that is, decreasing the entropy – of the value function^20^. This compression heightens the relative dominance of the best actions and thus facilitates exploitation and efficient exploration (Figure s1A-C). Critically, compression gives rise to entropy dynamics unpredicted by traditional RL (Figures 1G-H, s1D-E). These information dynamics scale with performance and non-verbal intelligence ^20^. To test the neural predictions of this model, we examine whether activity in the DAN is more consistent with an information-compressed value map or that from an otherwise identical traditional RL variant of SCEPTIC with long-term persistence of values (for details, see Methods).

Modern studies of functional brain connectivity in the human cortex reliably identify a dorsal attention network^33–36^, encompassing the temporo-occipital (putative human MT+), posterior parietal (IPS, SPL) and frontal premotor regions (FEF, PMv, PMd). A prominent parcellation of human functional network structure^33, 34^ further subdivides the DAN into two subnetworks along the caudo-rostral visuomotor gradient: the caudal subnetwork, consisting of MT+ and caudal PPC, and the rostral subnetwork consisting of rostral PPC and frontal premotor regions (Figure 2). Below, we describe activity across these four groups of regions. Since DAN subregions are characterized extensively in the macaque, for reference we label the human connectivity-based subregions according to their putative homology with monkey areas ^37–44^ (Figure s2, Table s1; “putative human” is omitted from region names below for simplicity since the interpretation of our results does not depend on precise homology between human and monkey areas).

**Figure 2.**
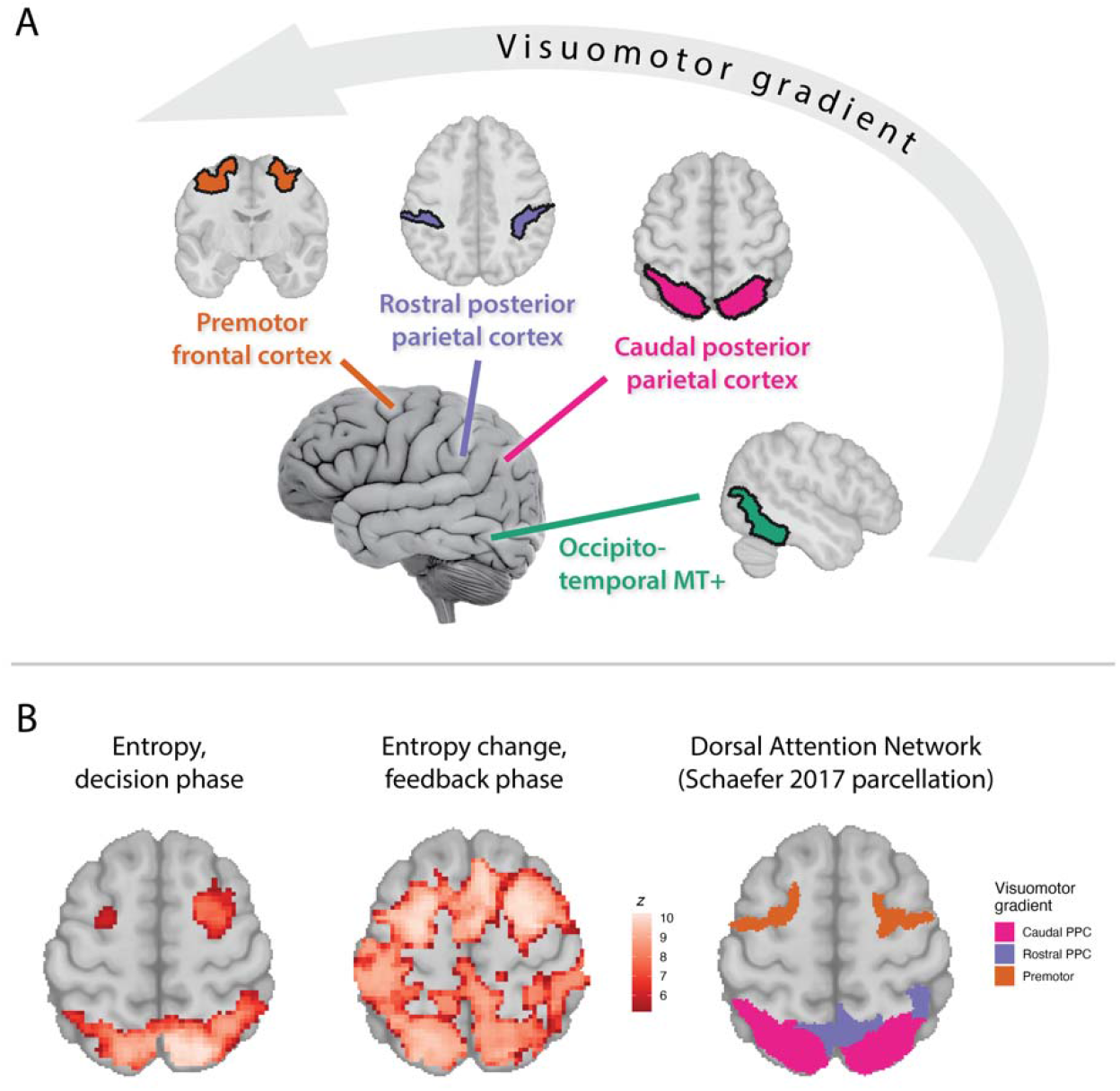
Human dorsal attention network (DAN) and responses to value entropy and its change. (A) DAN nodes arranged along the visuomotor transformation gradient, connectivity-based parcellation of Schaefer et al. (2018), details: Table s1. Detailed parcellation: Figure s2. (B) Responses to value entropy (left) and entropy change (middle), voxel-wise GLM; Schaefer DAN parcellation for the same axial slice (right; z = 55). Responses to reward/omission: Figure s3.

### Value information dynamics in the dorsal stream and the transition from exploration to exploitation

While fMRI and MEG cannot access option representations in individual neurons or subpopulations, our analyses of learned value information dynamics enabled us to adjudicate among competing accounts based on population-level measures. Specifically, if a subpopulation is recruited to represent the value of each option, the number of active subpopulations should scale with the information content of the learned value function. Thus, we can test whether maps contained in the human DAN undergo reinforcement-based updates as predicted by our model by regressing trial-by-trial entropy change against BOLD signal and posterior oscillatory power (Figure 1F). Entropy change reflects a global update to the dispersion of values for chosen and unchosen options and is distinct from prediction errors, which only reflect the local update to the chosen option, conceptually as well as statistically (|*r*|<0.1 for signed or absolute prediction errors across both samples reported here). Moreover, the information-compressing vs. traditional RL variants of the SCEPTIC model make different predictions about the nature of entropy change. Under the information-compressing model, entropy change has two components: 1) the decay of unchosen options, which reduces entropy of the value function and 2) value updates to the chosen action, which can increase or decrease entropy depending on whether the update promotes the dominant option relative to alternatives. By comparison, entropy change under the traditional RL model depends only on updates to the chosen action and entropy is generally higher relative to the information-compressing model ^20^. To demonstrate information-compressing learning dynamics, we contrasted the neural fit of our information-compressing model with an otherwise identical traditional RL comparator lacking information compression. We further ascertained that observations supporting it are not explained by random between-persons heterogeneity or confounds through fine-grained analyses of within-trial activity, stringent type I error control, sensitivity analyses, behavioral validation, and out-of-session and out-of-sample replication.

#### DAN BOLD scales with model-predicted value map information dynamics

Our whole-brain analysis revealed that the number of potentially advantageous options measured by model-predicted value entropy and its change (Figure 2B; Tables s2-s3) ^1^ recruited frontoparietal regions of the DAN, but not MT+, the basal ganglia or thalamus. Entropy change additionally recruited nodes of the cinguloopercular network (dorsal ACC and anterior insula/frontal operculum) and the rostrolateral prefrontal cortex.

#### Frontoparietal DAN nodes but not MT+ specifically track entropy change and not novelty

If DAN responses correlated with entropy change reflect value map updates, then activity should be modulated post-reinforcement. Indeed, analyses of within-trial BOLD activity revealed that entropy change modulated frontoparietal DAN activity post-outcome, particularly in caudal PPC and frontal-premotor nodes (Figure 3A). Responses in MT+ were much weaker, in contrast to responses to scalar value of the best option (Figure 3C), which were positive in MT+ and negative in fronto-parietal nodes. Effects of entropy change were evident with or without accounting for between-subject heterogeneity (individual random slopes), behavioral confounds (current and lagged response times) and spatially non-specific reinforcement features (scalar V_max_ [Figure 3C], reward/omission, prediction error all included as covariates in Figure 3A; model without covariates: Figure s4 A).

**Figure 3.**
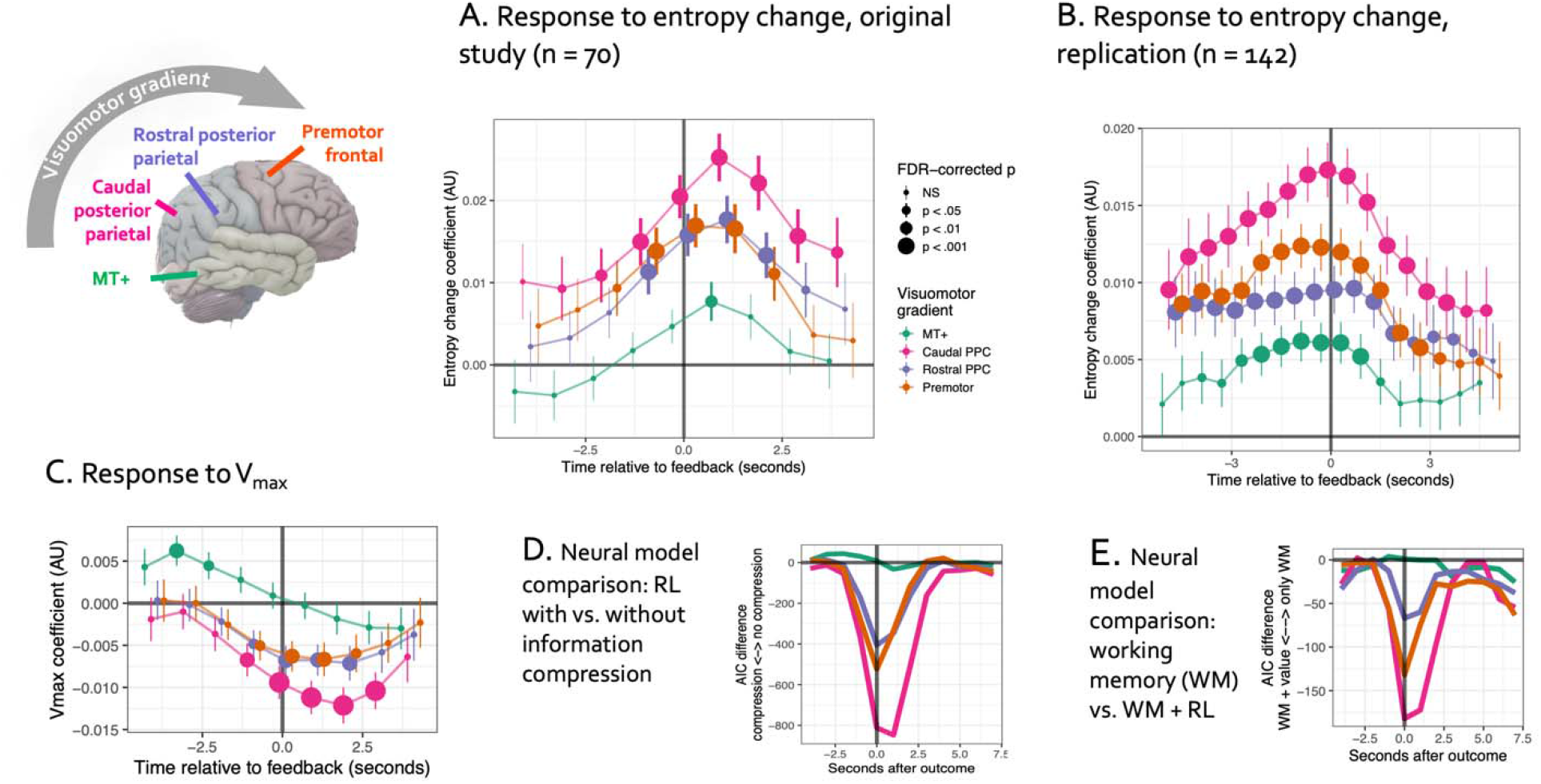
Information dynamics of the value function, DAN BOLD signal. (A) Responses to value entropy change (higher signal to increases reflecting a rising number of potentially valuable options), multi-level analysis of deconvolved BOLD signal from the original study, TR = 1s. (B) Same, replication sample of older adults with and without depression, TR = 0.6s. NB: Since BOLD response is smo oth, we can only interpret the peak response as indicating the timing of activity. (C) Response to scalar Vmax, original study. D. Neural model comparison providing evidence of information-compress ing rather than traditional reinforcement learning (RL). E. Analogous comparison demonstrating that DAN responses to entropy change cannot be explained solely in terms of spatial working memory updates.

Value entropy generally rises during early exploration when many options are sampled (Figure 1H), and one can argue that its association with neural responses is an artifact of novelty. To rule out this confound, we manipulated value entropy by changing the reward contingency every 40 trials without any explicit cues in a follow-up study of 142 older individuals with and without psychopathology. In this replication study, frontoparietal DAN responses to entropy change were qualitatively unchanged, even though we had excluded the first 10 trials from analyses to eliminate novelty effects (Figure 3B).

#### Frontoparietal BOLD dynamics specifically support reinforcement learning with information compression

A hallmark of traditional instrumental learning is the long-term persistence of option values. In contrast, in our information-compressing model of learning, values of preferred regions are maintained, whereas values of spatiotemporally distant alternatives decay. To find evidence of such compression, we compared the neural fit of entropy change signals from the information-compressing model vs. an otherwise identical model without compression. The information-compressing model better accounted for DAN responses to entropy change, particularly in PPC-caudal and frontal-premotor regions, but not in MT+ (AIC_selective_ – AIC_full_: ≥ -847, Figure 3D). Compression in the SCEPTIC model occurs as part of the reinforcement-driven update (Eq. 7), and as expected, its advantage peaks at reinforcement.

#### Information-compressing learning is not explained by working memory updates, but complements them

One alternative possibility is that participants perform the clock task by holding recent choices and outcomes in working memory and repeating recently rewarded choices. Human choices, however, are not adequately explained by such a process. The SCEPTIC-derived RT _Vmax_ explained substantial variance in choices even after accounting for a 5-trial buffer of choices and outcomes (fMRI session: t = 5.42, MEG session: t = 6.43,Table s4; outcomes >4 trials back had no detectable impact on choice).

Neural responses to the number of valuable options could also reflect updates to a spatial working memory buffer. Although the above behavioral analyses speak to the contrary, it was important to rule out this alternative account using neural data. By definition, spatial working memory contains the history of chosen locations (response times) and corresponding outcomes. Thus, to model working memory information content in a manner directly comparable with that of SCEPTIC, we encoded the selection history using the same representational structure (basis functions and spatial generalization gradient), adding a buffer of recent outcomes (detailed in Methods). We then predicted neural activity with the entropy and entropy change of the selection history and outcome history (reflecting working memory buffer updates, Figure s5 B-C) with and without corresponding SCEPTIC signals. As in our main analysis, responses to value entropy change peaked ∼1s post-outcome (Figure s5 A), and model fit improved by ≥-182 AIC points after adding SCEPTIC predictors (Figure 3E; and after accounting for random slopes of all entropy variables, ≥-1726 points). Overall, while our analyses replicate common findings of spatial working memory representations in the DAN and specifically PPC, they support parallel information-compressing updates of option values.

Another confound related to selection history is the potential impact on value entropy and its neural correlates of preceding RT swings, whether they reflect strategic exploration or stochastic or even off-task responses^45^. To rule out this possibility, we quantified a shift in the local distribution of choices as the summed Kullback-Leibler divergence (KLD; a metric of divergence between distributions) of response times for trials t-4, t-3, and t-2 from the local distribution of response times of the preceding three trials. Higher values of this measure reflect a history of larger RT swings. With (Figure 3A) or without (Figure s5A) this KLD measure as a covariate the entropy change effects were qualitatively unchanged, corroborating the notion that entropy change reflects value map updates rather than selection history. Interestingly, on trials following larger RT swings, we observed lower online rostral PPC activity and weaker frontoparietal responses to feedback (Figure s5B), potentially indicating lapses in sensorimotor activity and attention (see also a similar analysis of MEG below).

#### DAN sensitivity to the number of potentially valuable options predicts exploitation

The analyses above suggest that the DAN contains information-compressed value maps, but do these maps indeed govern the transition from exploration to exploitation? To answer this question, we tested whether individuals whose DAN activity better tracked with model-predicted value map updates (entropy change) made more value-sensitive, exploitative choices. We extracted entropy change regression coefficients (“betas”) for each DAN parcel from individual subjects’ whole-brain analyses and entered these as between-subjects predictors in a multilevel survival model (with timepoints nested within trials and trials nested within subjects) predicting the momentary rate (hazard) of response with SCEPTIC-derived within-trial momentary value and its interaction with the fMRI beta. This interaction was positive across the parcels, indicating that individual value sensitivity scaled with entropy change responses across the DAN. This effect was replicated out-of-session (Figure 4A; anatomical distribution of behavioral effect was preserved out of session, Figure 4D) and persisted in sensitivity analyses controlling for the non-decision time (censoring the first 1s of the interval) and avoidance of missing the response window (censoring the last 0.5s; Figure s6E-F). With individual random slopes, effects were similar in the original sample, surviving FDR correction in the out-of-session replication only in premotor parcels (Figure s6A-B). To ensure that this effect did not depend on the Cox model proportional hazards assumption, we tested it in an independent trial-level GLM predicting response times with SCEPTIC-derived *RT_Vmax_*, the location of the highest-valued option, which enabled us to account for additional behavioral confounds (Methods, fMRI Analyses) and between-subject heterogeneity in value sensitivity (random value slopes). The results were qualitatively unchanged (Figures 4B-C, s6).

**Figure 4.**
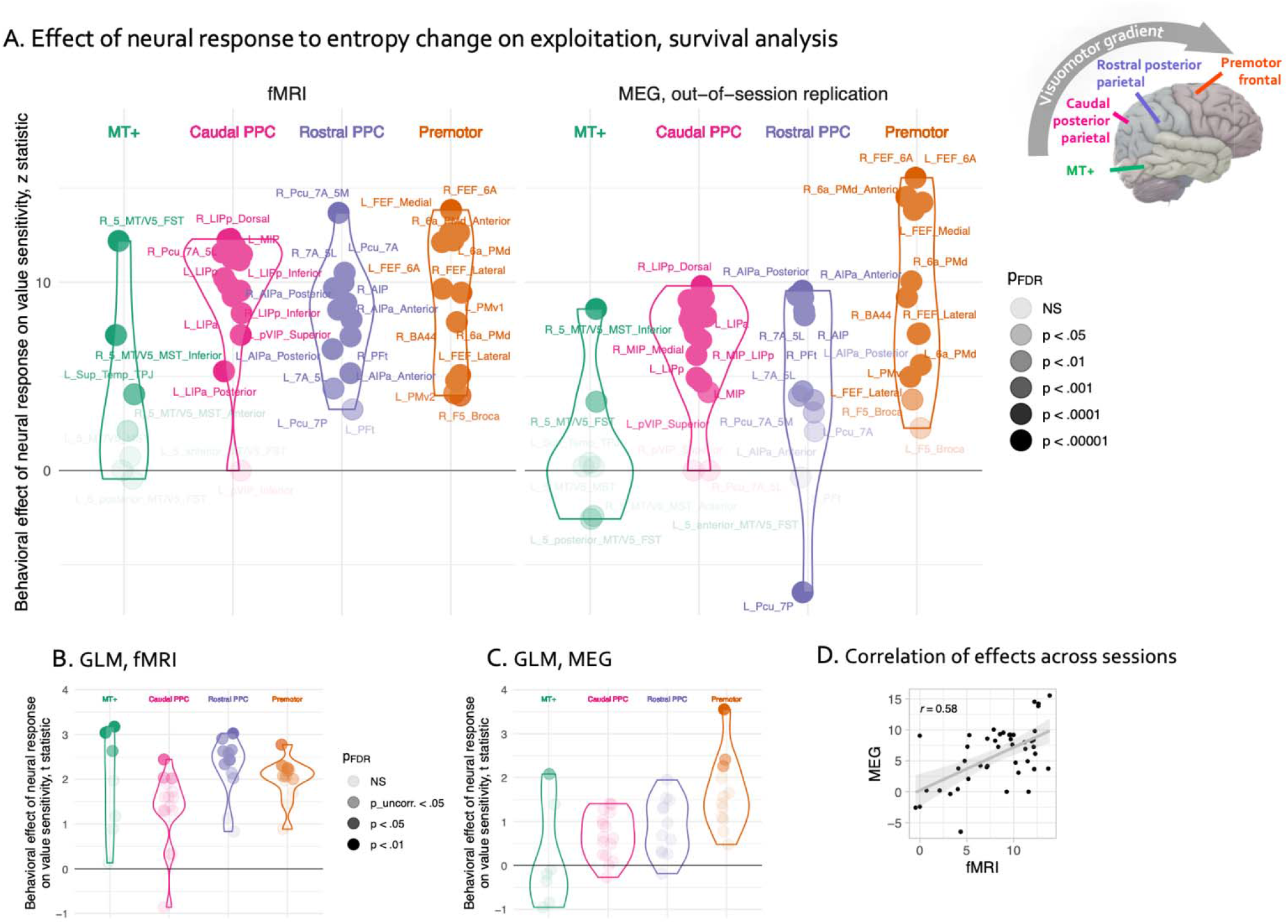
BOLD encoding of value information dynamics and behavioral exploitation. (A) Multi-level survival analyses examining how the individual’s neural response moderates their behavioral sensitivity to within-trial time-varying value. Left: original fMRI session. Right: replication, MEG session. Greater modulation of individual DAN BOLD response by value entropy change predicted more exploitative choices. (B), (C) Same, GLM analysis. (D) The anatomical pattern of brain-behavior associations was preserved across the original fMRI and replication sessions. Each dot represents a single DAN parcel as labeled in panel A.

#### Posterior β_1_/α suppression reflects value information dynamics

Having observed dynamic value maps encoded in fronto-parietal DAN BOLD, we sought to understand cortical oscillation dynamics that underlie them. A late (550-1000ms post-feedback) β_1_/α-band response has been reported on the clock task^46^, but its functional significance was unclear. We hypothesized that this response reflected an update to the parietal value map, with increases in the number of valuable options resulting in global desynchronization. Indeed, increases in entropy (and the number of valuable options) elicited suppression in the 7-17 Hz (β_1_/α) band at 400-750 ms, prominent in the posterior sensors (Figure 5 A, C). The reconstructed sources of this signal followed an anatomical distribution similar to the pattern observed in fMRI (Figure 5D vs. 5E). As in our analyses of BOLD, late β_1_/α suppression was not explained by behavioral confounds (reward, RT_t_, RT_t-1_, V_max_; Methods, MEG Analyses) and was strongly related to entropy change and absolute prediction errors, but not to reward/omission (Figure 5F), suggesting that β_1_/α oscillations encode updates to the entire map of chosen and unchosen options, with the chosen option commanding additional processing. This β_1_/α response was evident in two learnable and one unlearnable condition. However, it was almost abolished in CEVR (χ^2^(3) = 10.14, p < .018) where the probability/magnitude trade-off was the opposite of other conditions, indicating that the response was altered when outcomes did not match one’s expectations based on experience with previously encountered environments. This late suppression spread into the theta band, peaking at 600-800 ms and 3-6 Hz, evident mostly in posterior sensors. Additionally, an earlier burst of suppression at 8-17 Hz emerged immediately following response and ceased after the outcome (Figure 5A), suggesting that participants were at times anticipating an increase in global uncertainty based on their response.

**Figure 5.**
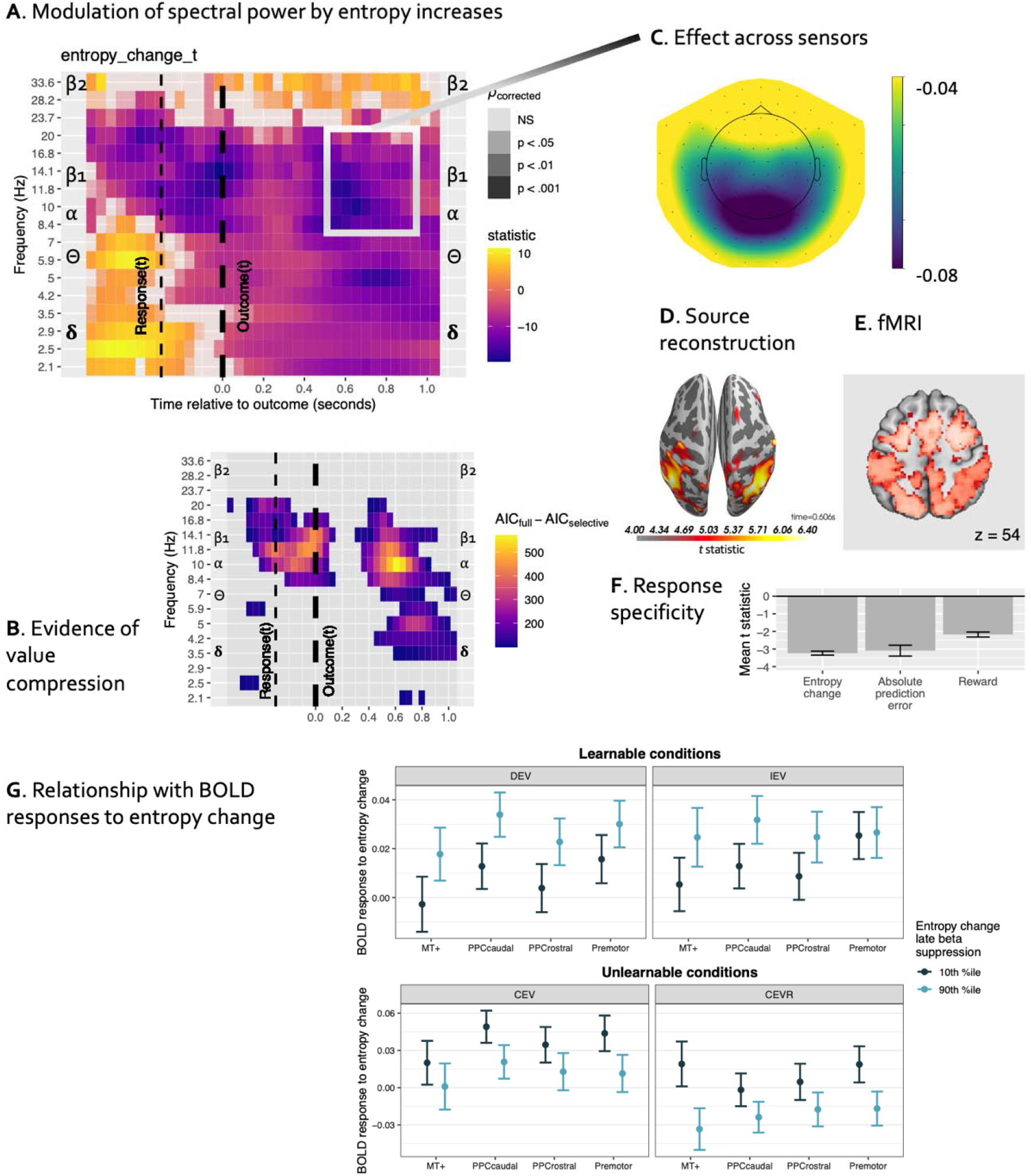
MEG: oscillatory responses to value information dynamics and exploitation, anatomical and functional relationship to BOLD signal. (A) Oscillatory response to value entropy change: cool colors represent de-synchronization to increases reflecting a ris ing number of potentially valuable options, most prominent between 7-17 Hz (β_1_/α) band at 400-750 ms. (B) Neural model comparison providing evidence of information-compressing rather than traditional RL (c.f. Figure 3D). Hot colors represent AIC difference favoring the information-compressing (selective maintenance) model. (C) β_1_/α de-synchronization was most evident in posterior sensors, consistent with a parietal source. (D) Source reconstruction localizes the β_1_/α suppression to the posterior parietal cortex. (E) fMRI BOLD map shown for comparison. (F) β_1_/α de-synchronization was much better explained by value entropy change or absolute reward prediction errors than by reward/omission, model controlling for individual random slopes. More negative t-statistics indicate a stronger effect. (G) Condition-level relationships between individuals’ BOLD and oscillatory responses to entropy change. Light blue bars represent individuals with stronger oscillatory responses (greater suppression to entropy increases). Y-axis: higher coefficient values indicate stronger BOLD response. X-axis: DAN regions from which BOLD response was extracted.

It is possible that, instead of β_1_/α desynchronization to entropy increases, our observations reflect β_1_/α synchronization to entropy *decreases* relative to baseline. We ruled out this possibility by separately examining the effects of entropy increase (vs. decrease or no change) and entropy decrease (vs. increase or no change; Figure s7A-B). Whereas entropy increases elicited massive suppression at 8-20 Hz peaking at 0.4-0.8s post-outcome and spreading into the theta band, entropy decreases did not elicit synchronization of a similar magnitude.

One could also argue that effects of entropy increases merely reflect a recent history of highly variable choices^45^ rather than updates to the distribution of learned value across competing options. Interestingly, while a recent history of RT swings measured by the Kullback-Leibler distance between RT _{t-3, t-2}_ and RT_t-1_, predicted suppression in the 7-16 Hz band (same model as above, Figure s7C), effects of entropy increases persisted while controlling for RT swings (Figure s7A), indicating that both selection and reinforcement history are encoded in β_1_/α oscillations.

Finally, manipulation effects are heterogeneous across individuals ^47^, and we verified that our findings were reliable after accounting for inter-individual heterogeneity by including the subject random slope of entropy change in our multi-level models (Figure s7D).

#### Posterior β_1_/α oscillation dynamics support information compression

Our analyses of BOLD indicated that reinforcement representations in the fronto-parietal DAN nodes were compressed as predicted by SCEPTIC. To understand whether similar information-compressing dynamics were reflected in oscillatory activity, we compared the fit of the MLM with entropy change regressor derived from either the information-compressing (exactly as in our main analysis above) or the traditional RL SCEPTIC model. Indeed, the information-compressing model dominated in the in the 8-17 Hz band at 400-750 ms, in the 3-6 Hz band at 600-800 ms and at 8-20 Hz peri-response (Figure 5 B), indicating that representations of competing options reflected in oscillatory activity displayed information-compressing dynamics predicted by the SCEPTIC model.

#### Posterior β_1_/α oscillation dynamics predict the explore-exploit transition

To understand whether β_1_/α suppression responses to an increased number of options (entropy change) scaled with exploitation we used models similar to those employed in fMRI analyses (Figure 4). To ensure that our results generalized across contingencies, we decomposed summary β_1_/α suppression responses into person-level means and condition-wise deviations. Person-level responses predicted exploitation (momentary value * b suppression response: *z* = 9.47, *χ*^2^(1) = 89.69, *p* < 10^-15^). This effect was robust to between-subject heterogeneity (random slope of value, fixed effect: z = 2.14, χ^2^(1) = 4.57, p = 0.0326) and replicated out-of-session (z = 13.03,χ^2^(1) = 169.88, p < 10^-15^), even after accounting for between-subject variability in the effect of value (z = 3.40, χ^2^(1) = 11.60,*p* < 0.001). After accounting for subject-level responses, no additional effect was observed, at the within-person, condition level (|z| ≤ 0.94,χ^2^(1) ≤ 0.86, p > 0.35), suggesting that the relationship of oscillatory responses and behavioral exploitation manifests at the between-person level.

#### Magnitude of posteriorβ_1_/α response scales negatively with BOLD, but only in learnable conditions

Condition-level β_1_/α synchrony scaled negatively with BOLD responses across DAN, but only in learnable conditions, while the opposite pattern was seen in unlearnable conditions (Figure 5G, β_1_/α response main effect: χ^2^(1) = 10.65, p = 0.0011,β_1_/α response *condition χ^2^(3) = 36.49, p < 10^-7^), suggesting that β_1_/α suppression and/or BOLD are differentially sensitive to the presence of an objective value maximum, and potentially also to the match between current and previously encountered contingencies.

In summary, we found that the fronto-parietal nodes of the dorsal stream represented a compressed reinforcement history, mapping values of potentially valuable options as did posterior β_1_/α oscillations. These neural dynamics predicted a successful transition from exploration to exploitation.

## Discussion

When exploring and exploiting a few discrete options, primates rely on choice and reinforcement histories encoded in the striatum and amygdala. Additional demands, however, arise when moving rapidly through space and choosing when to harvest a reward. Our multimodal imaging study of human BOLD signal and cortical oscillations revealed that reward-based learning in such an environment involves dynamic value maps in the dorsal stream. More specifically, we found that BOLD signals in the PPC and premotor cortex increased and posterior β_1_/α oscillations desynchronized in response to increases in the number of valuable options and, correspondingly, uncertainty about the best option. This global uncertainty was quantified by changes in the entropy of the learned value function captured by our computational model. These BOLD and oscillatory dynamics predicted a successful behavioral transition from exploration to exploitation, with out-of-session and out-of-sample replication. BOLD dynamics consistent with map updates were seen throughout the parietal and frontal nodes of the dorsal stream, but not in the occipito-temporal MT+.

Much debate about maps in the dorsal stream, especially in the PPC, has focused on what they represent. They have been suggested to encode attentional priority ^48–51^, the intention to move^52^, expected value vs. salience of stimuli^25, 26, 53^, or expected information gain^54, 55^. Disagreements between studies are not entirely explained by anatomical heterogeneity or methodological differences ^10^. Rather, this debate may be resolved in part by recognizing that real-world motor programs are inextricable from visuospatial and value-laden representations of targets. This aligns with the affordance perspective in which sensorimotor systems continuously encode opportunities for action emerging in the immediate environment ^9, 56^. Affordance representations throughout the dorsal stream multiplex visual, oculomotor, motor, and somatosensory information ^57–59^. We observed that the nodes in the dorsal stream along the visuomotor gradient from caudal PPC to premotor cortex respond similarly to the values of options, consistent with the notion of pragmatic, multimodal affordance representations. These dynamics were considerably weaker in the MT+, which instead responded to the recent reward and long-term value, potentially indicating that processing of visual motion (here, ball motion around the clock face) was enhanced when the expected reward rate was high, as electrophysiological studies suggest ^60^. Thus, fronto-parietal regions but not MT+ contain a spatially structured value map for all options.

Our results address the critical question of how competition between multiple affordances is resolved^9, 61, 62^. Cisek and colleagues speculate that the prevailing affordance in output regions is determined by both reinforcement learning and goals signaled to the dorsal stream by ventral prefrontal systems^63–65^. We find that chosen and unchosen option values are updated on the PPC map within 0.4–0.7s of reinforcement, predicting exploitation of the highest-valued option. Crucially, dorsal stream value map updates and behavior were better described by an information-compressing reinforcement learning algorithm relative to traditional instrumental learning, even after accounting for updates to the visuospatial WM buffer^66^. This algorithm selectively maintains preferred actions, compressing information about learned values and supporting more efficient exploitation during continuous visuomotor interactions.

Our observations are not easily explained by an earlier account of reward learning in discrete spaces postulating that it relies on early (0.2-0.4s) traditional RL updates and later (0.4-0.7s) working memory updates^67^. During continuous sensorimotor interaction we observe both value map and spatial working memory buffer updates 0.4-0.7s post-outcome. How does the PPC obtain information about incoming reinforcement? The identity of a reward or goal state may be cached in the dorsal and ventral stream^65^. Thus, upon reaching the goal, reinforcement may be almost instantaneous ^63^. Reinforcement may also be signaled meso-striato-thalamo-cortical reward prediction errors. However, these signals are unlikely to arrive early enough to enable the observed value map updates^63^ and thus may shape learning only at slower timescales. In any case, since updates to values of *unchosen* regions of the continuous space must account for the spatial proximity of the sampled point, they can only be updated by a network that contains a full map, such as the PPC. Our results suggest that a dual-systems account of competing frontoparietal WM and meso-striatal RL controllers ^67^ may not extend to the rapid continuous sensorimotor interaction. Affordance competition must in part be resolved through reinforcement learning in the dorsal stream, with information compression facilitating a “within-system” decision ^68^.

The connection between information-compressing RL and β_1_/α oscillations bridges our population-level account of option competition in the dorsal stream with biophysically realistic circuit models. Gelastopoulos, Whittington and Kopell ^18^ propose that competing representations in the parietal cortex are carried by β_1_/α-synchronized ensembles organized along cortical columns. It is thought that recurrent excitation stabilizes preferred options, while lateral inhibition suppresses non-preferred alternatives^18, 69, 70^. In line with these circuit-level models and empirical studies, we propose that during the value-guided exploration of the sensorimotor space, recruitment of many ensembles produces an asynchronous oscillatory output, reflecting greater entropy of the value map and global uncertainty about the best action (Figure 6). When an option is preferentially sampled and reinforced, the regional output becomes dominated by the β_1_/α -synchronized ensemble representing this option, promoting exploitation. Compression of the value function may depend on lateral inhibition in which dominant ensembles suppress and even highjack the β_1_/α output of competing counterparts, reducing their likelihood of behavioral selection^18, 69^.

**Figure 6.**
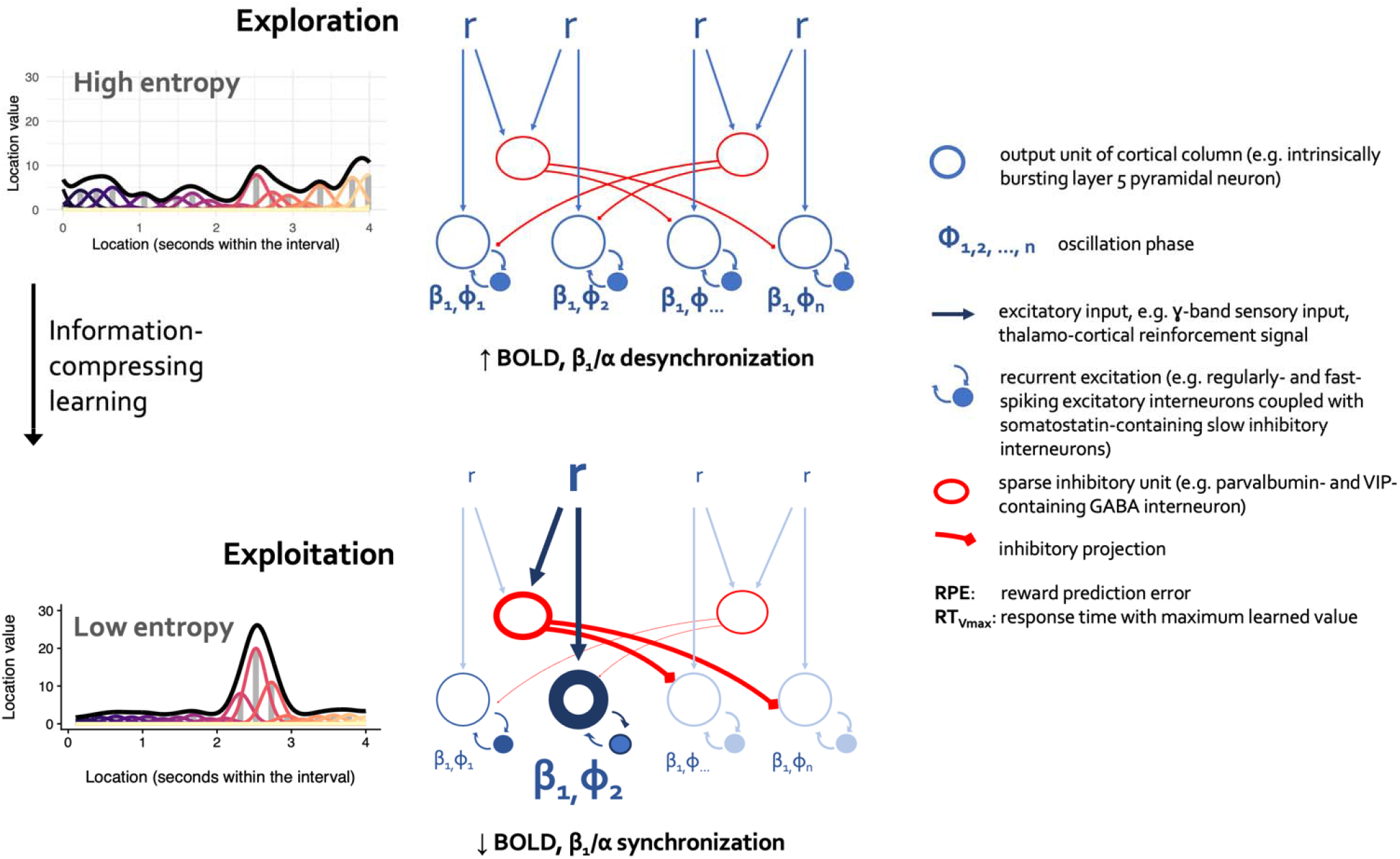
Option competition in the posterior parietal cortex: conceptual model. Top: exploration mode. Top left: Multiple options compete for selection and the entropy of the value function is high. Top right: Competing options are represented by neuronal subpopulations with each producing oscillatory output with a distinct phase. Bottom: exploitation mode. Top left: following information-compressing learning, a global value maximum emerges. Top right: as a result of recurrent excitation and lateral inhibition, the subpopulation representing the dominant option begins to dominate the output. After Gelastopoulos, Whittington and Koppel (2019; biophysical model of β_1_/α-stabilized competing cortical populations) and Mysore and Kothari (2020; computational models of competitive selection).

We observe a clear functional correspondence between dorsal stream BOLD and posterior β_1_/α desynchronization: value entropy change modulated BOLD positively (Figure 3A-B, 4B-c) and β_1_/α power negatively (Figure 5A, E). Interestingly, BOLD and β_1_/α responses were correlated only in learnable contingencies. Moreover, oscillations were sensitive to a mismatch with previous experience: in a contingency with an unexpectedly reversed probability/magnitude tradeoff, oscillatory responses no longer tracked entropy dynamics. While the spatial resolution of fMRI complements the temporal resolution of electrophysiology, BOLD and oscillatory power capture different aspects of cortical local field potentials. Their empirical correlations are generally positive in gamma band and negative in α and β (which contain additional unique information about BOLD), with β but not α suppression accelerating increases and delaying decreases in BOLD^71–73^. Thus, our fMRI and MEG findings align and ostensibly reflect updates to the dynamic value map. These functional properties were not shared by other frequency bands such as high beta, theta and delta.

The strengths of our study include consistent findings of value entropy dynamics in the human dorsal stream across MEG and fMRI modalities, across sessions, and in a separate fMRI sample. Out-of-session and out-of-sample replications increase confidence in the observed links between dynamic maps and behavioral exploration/exploitation. Our computational model comparisons supported the conclusion that value maps in the dorsal stream are likely shaped by an information-compressing RL process that cannot be explained by traditional instrumental learning or WM buffer accounts. Experimental manipulation of reinforcement in a continuous space and our RL model provided access to a spatially structured value vector, dissociating global from local updates. Our novel multilevel analyses revealed parallel within-trial temporal dynamics of cortical oscillations and deconvolved BOLD signals. Finally, our observations of value entropy dynamics replicated in a modified experiment with unsignaled reversals, ruling out novelty as an alternative explanation.

The main limitation of our study is the lack of a causal manipulation that would isolate contributions of various DAN nodes to resolving the explore-exploit dilemma, although our findings of dynamic value maps in the dorsal stream are in line with human transcranial stimulation studies^53, 74^. Human neural stimulation and rodent optogenetic studies are needed to test the model of value-dependent affordance competition articulated here. It is also difficult to know to what extent our findings generalize to non-temporal spaces and to punishments as opposed to rewards. Finally, MEG and fMRI data were collected in separate sessions, enabling out-of-session replication, but precluding an analysis of simultaneous BOLD and cortical oscillation recordings.

In conclusion, exploration and exploitation of a continuous sensorimotor space depend on dynamic value maps in the dorsal stream, particularly in the PPC. Our observations suggest that the dorsal stream selectively maintains values of preferred options and compresses out inferior, spatiotemporally distant alternatives. Indeed, as circuit models of option competition in the dorsal cortex suggest, β_1_/α oscillatory output of posterior cortical subpopulations desynchronizes when more competing options emerge and synchronizes when non-preferred alternatives are compressed out. Compression notwithstanding, we show that option values in PPC persist beyond the timescale of the WM buffer. Our results support the affordance competition view of maps in dorsal cortex and are at odds with the notion that sensorimotor choices require a sequence of temporally distinct sensory, reward, cognitive and motor computations. Altogether, our study sheds light on how primates, including humans, track the values of alternative options in complex, rapidly changing environments.

## Methods

### Participants

Participants in the original study were 70 typically developing adolescents and young adults aged 14– 30 (*M* = 21.4, *SD* = 5.1). Thirty-seven (52.8%) participants were female and 33 were male. Prior to enrollment, participants were interviewed to verify that they had no history of neurological disorder, brain injury, pervasive developmental disorder, or psychiatric disorder (in self or first-degree relatives). Participants in the replication study were 143 middle-aged and older adults aged 50-80 (*M* = 62.2, *SD* = 6.8), 80 (56%) were female and 62 were male; 101 were diagnosed with DSM-IV non-psychotic major depression. Individuals with a history of psychosis, mania, neurological conditions of the brain and current substance use disorders were excluded from the replication study. Participants and/or their legal guardians provided informed consent or assent prior to participation in both studies. Experimental procedures for this study complied with Code of Ethics of the World Medical Association (1964 Declaration of Helsinki) and the Institutional Review Board at the University of Pittsburgh (protocols PRO10090478 and STUDY19030288). Participants were compensated $75 for completing the original experiment and $150 for completing the replication study, which included other experiments.

### Procedure

#### Original study

As part of a larger study, participants completed an exploration and learning task (“clock task”; Figure 1A, and detailed below) in separate magnetoencephalography (MEG) and functional MRI (fMRI) sessions. The order of the fMRI and MEG sessions was counterbalanced (fMRI first *n* = 34, MEG first *n* = 36) and the sessions were separated by 3.71 weeks on average (SD = 1.59 weeks).

During the fMRI session, participants completed eight runs of the clock task (based on Moustafa et al., 2008). Runs consisted of 50 trials in which a green dot revolved 360° around a central stimulus over the course of 4s. Participants pressed a button to stop the dot, which ended the trial. They then received a probabilistic reward for the chosen response time (RT) according to one of four time-varying contingencies, two learnable (increasing and decreasing expected value) and two unlearnable. All contingencies were monotonic but featured reward probability/magnitude tradeoffs that made learning difficult (see^20^ for more detailed analyses of the task). After each response, participants saw the probabilistic reward feedback for 0.9s. If participants failed to response within 4s, they received zero points.

The central stimulus was a face with a happy expression or fearful expression, or a phase-scrambled version of face images intended to produce an abstract visual stimulus with equal luminance and coloration. Faces were selected from the NimStim database ^75^. All four contingencies were collected with scrambled images, whereas only IEV and DEV were also collected with happy and fearful faces. The effects of the emotion manipulation will be reported in a separate manuscript because they are not central for the examination of the neural substrates of exploration and exploitation on this task.

Each trial was followed by an intertrial interval (ITI) that varied in length according to an exponential distribution. To maximize fMRI detection power, the sequence and distribution ITIs were derived using a Monte Carlo approach implemented by the *optseq2* command in *FreeSurfer* 5.3. More specifically, we simulated five million possible ITI sequences consisting of 50 trials each and retained the top 320 orders based on their estimation efficiency. For each subject, the experiment software randomly sampled 8 of these efficient ITI sequences, which were used for the durations of ITIs in the task.

During the MEG session, participants completed eight runs of the same task. The contingencies and trial structure were identical to fMRI (see Figure 1A), requiring participants to respond within a four-second interval to maximize the points they earned. Given the lower signal-to-noise ratio of MEG relative to fMRI, runs consisted of 63 trials each.

As detailed in the results, the behavioral data from the MEG and fMRI sessions were used to test the out-of-session consistency of brain-behavior effects identified by each modality. This enabled us to establish whether individual differences in dorsal stream activity and exploration/exploitation represented stable tendencies vs. patterns incidental to a single experimental session.

#### Replication study

Procedures of the replication study were similar, but no MEG data were collected. Participants completed 240 trials of the clock task in two runs. Only IEV and DEV contingencies were employed. To dissociate value entropy from novelty, the contingency reversed every 40 trials unbeknownst to the participants. Trials were extended to 5s to accommodate slower psychomotor speed in this older sample.

### Imaging Acquisition and Processing Methods

#### fMRI acquisition

Neuroimaging data during the clock task were acquired in a Siemens Tim Trio 3T scanner for the original study and Siemens Tim Prisma 3T scanner for the replication study at the Magnetic Resonance Research Center, University of Pittsburgh. Due participant-dependent variation in response times on the task, each fMRI run varied in length from 3.15 to 5.87 minutes *M*(= 4.57 minutes, *SD* = 0.52). Functional imaging data for the original/replication study were acquired using a simultaneous multislice sequence sensitive to BOLD contrast, TR = 1.0/0.6s, TE = 30/27ms, flip angle = 55/45°, multiband acceleration factor = 5/5, voxel size = 2.3/3.1mm^3^. We also obtained a sagittal MPRAGE T1-weighted scan, voxel size = 1/1mm ^3^, TR = 2.2/2.3s, TE = 3.58/3.35ms, GRAPPA 2/2x acceleration. The anatomical scan was used for coregistration and nonlinear transformation to functional and stereotaxic templates. We also acquired gradient echo fieldmap images (TEs = 4.93/4.47ms and 7.39/6.93ms) for each subject to mitigate inhomogeneity-related distortions in the functional MRI data.

#### Preprocessing of fMRI data

Anatomical scans were registered to the MNI152 template^76^ using both affine (ANTS SyN) and nonlinear (FSL FNIRT) transformations. Functional images were preprocessed using tools from NiPy ^77^, AFNI (version 19.0.26)^78^, and the FMRIB software library (FSL version 6.0.1) ^79^. First, slice timing and motion coregistration were performed simultaneously using a four-dimensional registration algorithm implemented in NiPy ^80^. Non-brain voxels were removed from functional images by masking voxels with low intensity and by the *ROBEX* brain extraction algorithm^81^. We reduced distortion due to susceptibility artifacts using fieldmap correction implemented in FSL FUGUE.

Participants’ functional images were aligned to their anatomical scan using the white matter segmentation of each image and a boundary-based registration algorithm ^82^, augmented by fieldmap unwarping coefficients. Given the low contrast between gray and white matter in echoplanar scans with fast repetition times, we first aligned functional scans to a single-band fMRI reference image with better contrast. The reference image was acquired using the same scanning parameters, but without multiband acceleration. Functional scans were then warped into MNI152 template space (2.3mm output resolution) in one step using the concatenation of functional-reference, fieldmap unwarping, reference-structural, and structural-MNI152 transforms. Images were spatially smoothed using a 5mm full-width at half maximum (FWHM) kernel using a nonlinear smoother implemented in FSL SUSAN. To reduce head motion artifacts, we then conducted an independent component analysis for each run using FSL MELODIC. The spatiotemporal components were then passed to a classification algorithm, ICA-AROMA, validated to identify and remove motion-related artifacts ^83^. Components identified as noise were regressed out of the data using FSL regfilt (non-aggressive regression approach). ICA-AROMA has performed very well in head-to-head comparisons of alternative strategies for reducing head motion artifacts^84^. We then applied a .008 Hz temporal high-pass filter to remove slow-frequency signal changes ^85^; the same filter was applied to all regressors in GLM analyses. Finally, we renormalized each voxel time series to have a mean of 100 to provide similar scaling of voxelwise regression coefficients across runs and participants.

#### Treatment of head motion

In addition to mitigating head motion-related artifacts using ICA-AROMA, we excluded runs in which more than 10% of volumes had a framewise displacement (FD) of 0.9mm or greater, as well as runs in which head movement exceeded 5mm at any point in the acquisition. This led to the exclusion of 11 runs total, yielding 549 total usable runs across participants. Furthermore, in voxelwise GLMs, we included the mean time series from deep cerebral white matter and the ventricles, as well as first derivatives of these signals, as confound regressors ^84^.

#### MEG Data acquisition

MEG data were acquired using an Elekta Neuromag VectorView MEG system (Elekta Oy, Helsinki, Finland) in a three-layer magnetically shielded room. The system comprised of 306 sensors, with 204 planar gradiometers and 102 magnetometers. In this project we only included data from the gradiometers, as data from magnetometers added noise and had a different amplitude scale. MEG data were recorded continuously with a sampling rate of 1000 Hz. We measured head position relative to the MEG sensors throughout the recording period using 4 continuous head position indicators (cHPI) that continuously emit sinusoidal signals, and head movements were corrected offline during preprocessing. To monitor saccades and eye blinks, we used two bipolar electrode pairs to record vertical and horizontal electrooculogram (EOG).

#### Preprocessing of MEG data

Flat or noisy channels were identified with manual inspections, and all data preprocessed using the temporal signal space separation (TSSS) method ^86, 87^. TSSS suppresses environmental artifacts from outside the MEG helmet and performs head movement correction by aligning sensor-level data to a common reference^88^. This realignment allowed sensor-level data to be pooled across subjects group analyses of sensor-space data. Cardiac and ocular artifacts were then removed using an independent component analysis by decomposing MEG sensor data into independent components (ICs) using the infomax algorithm ^89^. Each IC was then correlated with ECG and EOG recordings, and an IC was designated as an artifact if the absolute value of the correlation was at least three standard deviations higher than the mean of all correlations. The non-artifact ICs were projected back to the sensor space to reconstruct the signals for analysis. After preprocessing, data were epoched to the onset of feedback, with a window from -0.7 to 1.0 seconds. Trials with gradiometer peak-to-peak amplitudes exceeded 3000 fT/cm were excluded. For each sensor, we computed the time-frequency decomposition of activity on each trial by convolving time-domain signals with Morlet wavelet, stepping from 2 to 40 Hz in logarithmic scale using 6 wavelet cycles. This yielded trial-level time-frequency data that were amenable to multilevel models across frequencies and peri-feedback times.

### Computational Model of Behavior

#### Core architecture of SCEPTIC reinforcement learning (RL) model

The SCEPTIC model represents the one-dimensional space/time of the clock task using a set of unnormalized Gaussian radial basis functions (RBFs) spaced evenly over an interval *T* in which each function has a temporal receptive field with a mean and variance defining its point of maximal sensitivity and the range of times to which it is sensitive, respectively (a conceptual depiction of the model is provided in Figure 1). The primary quantity tracked by the basis is the expected value of a given choice (response time; we use the intuitive term *value* for continuity with prior studies of PPC maps ^25, 26, 53^, however since this estimate does not converge on the true reward rate, it is technically a *preference*, see text following eq. 7). To represent time-varying value, the heights of the basis functions are scaled according to a set of *B* weights, **w** = [w_1_, w_2_,…, w_b_]. The contribution of each basis function to the integrated value representation depends on its temporal receptive field:

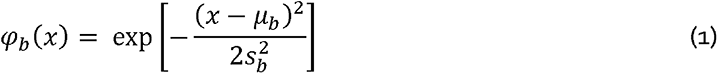

where *x* is an arbitrary point within the time interval *T*, µ_b_ is the center (mean) of the RBF and s^2^ is its variance. And more generally, the temporally varying expected value function on a trial *i* is obtained by the multiplication of the weights with the basis:

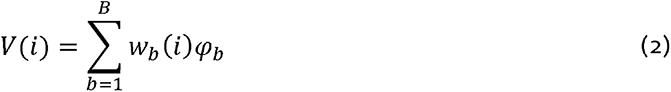

For the clock task, where the probability and magnitude of rewards varied over the course of four-second trials, we spaced the centers of 24 Gaussian RBFs evenly across the discrete interval and chose a fixed width, 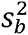, to define the temporal variance (width) of each basis function. More specifically, 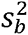 was chosen such that the distribution of adjacent RBFs overlapped by approximately 50% (for details and consideration of alternatives, see^20^).

The basic model, referred to as *traditional RL* in Results, learns the expected values of different response times by updating each basis function *b* according to the equation:

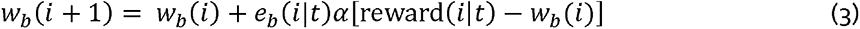

Where *i* is the current trial in the task, *t* is the observed response time (aka RT), and reward(i|t) is the reinforcement obtained on trial *i* given the choice *t*. Prediction error updates are weighted by the learning rate a and the temporal generalization function or eligibility *e*. To avoid tracking separate value estimates for each possible moment, feedback obtained at a given response time *t* is propagated to adjacent times. Thus, to implement temporal generalization of expected value updates, we used a Gaussian RBF centered on the response time *t,* having width 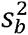. The eligibility *e_b_* of a basis function *φ_b_* to be updated by prediction error is defined as its overlap with the temporal generalization function, *g*:

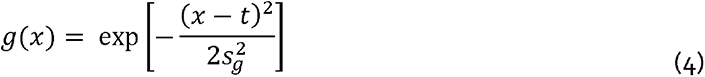

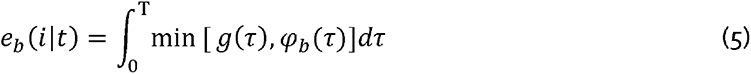

where τ represents an arbitrary timepoint along the interval *T*. Thus, for each RBF *b*, a scalar eligibility e_b_ between zero and one represents the proportion of overlap between the temporal generalization function and the receptive field of the RBF ^90^. In the case of complete overlap, where the response time is perfectly centered on a given basis function, *e_b_* will reach unity, resulting a maximal weight update according to the learning rule above. Conversely, if there is no overlap between an RBF and the temporal generalization function, *e_b_* will be zero and that RBF will receive no update. Importantly, for the eligibility to be bounded on interval [0,1], the basis functions are each normalized to have an area under the curve of unity (i.e., representing probability density). Here, we also fixed the width of the generalization function to match the basis (i.e., 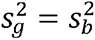).

The SCEPTIC model selects an action based on a softmax choice rule, analogous to simpler reinforcement learning problems (e.g., two-armed bandit tasks ^91^). For computational speed, we arbitrarily discretized the interval into 100ms time bins such that the agent selected among 40 potential responses (i.e., a multinomial representation). At trial i the agent chooses a response time in proportion to its expected value:

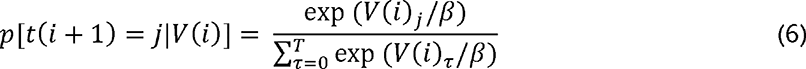

where *j* is a specific response time and the temperature parameter *β* controls value sensitivity such that choices became more stochastic and less value-sensitive at higher *β* values.

#### Information-compressing RL with selective maintenance

Importantly, as detailed previously ^20^, a model that selectively maintained frequently chosen, preferred actions far outperformed model alternative models. Specifically, basis weights corresponding to non-preferred, spatiotemporally distant actions revert toward a prior in inverse proportion to the temporal generalization function:

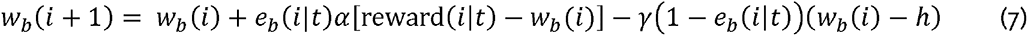

where γ is a selective maintenance parameter between zero and one that scales the degree of reversion toward a point *h*, which is taken to be zero here for parsimony, but could be replaced with an alternative prior expectation. Our primary fMRI analyses used signals derived from fitting the information-compressing RL model (eq. 7) to participants’ behavior, while comparisons with traditional RL used the model with the learning rule described in equation 3. Two features supported by computational studies and tests against human behavior ^20^, (i) the decay in the weights of unchosen alternatives (eq. 7) and (ii) calculation of prediction errors based on individual element weightsw_b_ (eq. 3, 7) rather than the total value estimate *V*(*i*) (eq. 2) preclude w_b_ or *V*(*i*) from converging on the true reward rate. While we refer to *w_b_* as *expected values* for continuity with previous studies of the parietal cortex^25, 26, 53^, *w_b_* are closer to *preferences*in policy gradient algorithms ^22, 92, 93^. Here, we focus on testing the hypothesis of information compression (eq. 8) and make no strong claims about whether representations of reinforcement in PPC constitute expected values or preferences. Although taken together with our earlier behavioral and computational results, neural model comparisons reported here can be taken to favor the preferences hypothesis, a definitive adjudication will require new experiments. Value vs. policy learning accounts are not necessarily mutually exclusive since actor-critic algorithms combine both approaches.

As detailed in the Results, we defined the information content of the learned value distribution as Shannon’s entropy of the normalized basis weights (the trial index *i* is omitted for simplicity of notation):

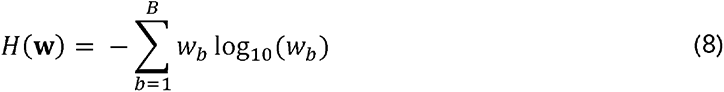

We further sought to examine whether entropy responses in the dorsal attention network were consistent with the information-compressing selective maintenance model. To test the specificity of the representation, we conducted analyses using entropy calculated from the information-compressing SCEPTIC selective maintenance model (equation 7) vs. entropy from a traditional RL, full-maintenance counterpart that did not decay the values of unchosen actions (equation 3; detailed model comparisons provided in^20^).

#### Working memory model

We argue that information dynamics attributable to value entropy from the SCEPTIC information-compressing model best explain the exploration-exploitation transition in behavior and updates to the value map in the DAN. Yet, one could imagine that that the DAN relies solely on a spatial working memory representation with a buffer containing locations and outcomes of recent choices. In turn, the information content of this buffer might be sufficient to explain DAN BOLD activity.

We first examined whether a WM process alone is sufficient to explain human choices without invoking information-compressing RL. We used a multi-level regression model as described below (*Brain-behavior fMRI analyses using regression coefficients from model-based fMRI GLM analyse*) *s*to predict the participant’s current RT with *k* preceding RTs representing the selection history buffer and their interactions with reward/omission representing the outcome buffer. This analysis revealed no effect of outcomes beyond 4 trials (Supplementary Table 4).To assess the effect of the more remote reinforcement history not captured by the last *k* choices and outcomes, we then tested the incremental contribution of the RT_Vmax_ (time of the learned value maximum) derived from the SCEPTIC model.

To conduct a conclusive test of this alternative account, we created a strong working memory-only comparator model that adopted the SCEPTIC RBF representation, using the basis weights to store the buffer of recently chosen locations, alongside a separate vector of recent outcomes. Specifically, the model remembered the past *k* choices by placing a unit-height eligibility functions at these locations and taking the sum, forming a selection history function, s(x). *k* was empirically estimated as 4 using multi-level linear regression (Supplemental Table 4). In turn, this representation of selection history was combined with the RBF by multiplying the selection history function and basis, yielding working memory basis weights 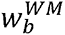 whose height scaled with the selection history.

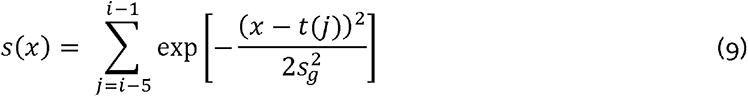

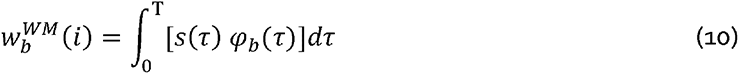

where *i* represents the current trial and *t(j)* is the response time on the *i*th previous trial. In turn, entropy can be calculated on the selection history basis weights in the same fashion as in the regular SCEPTIC model (Equation 8). Outcome history was simply represented by a vector o containing 1s for rewards and 0s for reward omissions in the past four trials. This implementation did not require estimating free parameters from behavior. Thus, total entropy or information content of working memory *H^WM^* is the joint entropy of the selection and outcome buffers:

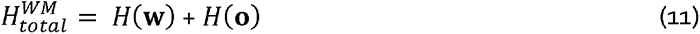

#### Fitting of SCEPTIC model to behavioral data

SCEPTIC model parameters were fitted to individual choices using an empirical Bayesian version of the Variational Bayesian Approach ^94^. The empirical Bayes approach relied on a mixed-effects model in which individual-level parameters were assumed to be sampled from a normally distributed population. The group’s summary statistics, in turn, were inferred from individual-level posterior parameter estimates using an iterative variational Bayes algorithm that alternates between estimating the population parameters and the individual subject parameters. Over algorithm iterations, individual-level priors are shrunk toward the inferred parent population distribution, as in standard multilevel regression. Furthermore, to reduce the possibility that individual differences in voxelwise estimates from model-based fMRI analyses reflected differences in the scaling of SCEPTIC parameters, we refit the SCEPTIC model to participant data at the group mean parameter values. This approach supports comparisons of regression coefficients across subjects and reduces the confounding of brain-behavior analyses by the individual fits of the computational model to a participant’s behavior. We note, however, that our results were qualitatively the same when model parameters were free to vary across people (additional details available from the corresponding author upon request).

### fMRI analyses

#### Voxelwise fMRI general linear model analyses

Voxelwise general linear model (GLM) analyses of fMRI data were performed using FSL version 6.0.4^79^. Single-run analyses were conducted using FSL FILM, which implements an enhanced version of the GLM that corrects for temporal autocorrelation by prewhitening voxelwise time series and regressors in the design matrix^85^. For each design effect, we convolved a duration-modulated unit-height boxcar regressor with a canonical double-gamma hemodynamic response function (HRF) to yield the model-predicted BOLD response. All models included convolved regressors for the clock and feedback phases of the task.

Moreover, GLM analyses included parametric regressors derived from SCEPTIC. For each whole-brain analysis, we added a single model-based regressor from SCEPTIC alongside the clock and feedback regressors. Results were qualitatively unchanged, however, when all SCEPTIC signals were included as simultaneous predictors, given the relatively low correlation among these signals. For each model-based regressor, the SCEPTIC-derived signal was mean-centered prior to convolution with the HRF. The reward prediction error and entropy change signals were aligned with the feedback, whereas entropy was aligned with the clock (decision) phase. Furthermore, for regressors aligned with the clock phase, which varied in duration, we sought to unconfound the height of the predicted BOLD response due to decision time from the parametric influence of the SCEPTIC signal. Toward this end, for each trial, we convolved a duration-modulated boxcar with the HRF, renormalized the peak to unity, multiplied the regressor by the SCEPTIC signal on that trial, then summed across trials to derive a single model-based regressor (cf. processing time versus intensity of activation in ^95^).

Parameter estimates from each run were combined using a weighted fixed effects model in FEAT that propagated error variances from the individual runs. The contrasts from the second-level analyses were then analyzed at the group level using a mixed effects approach implemented in FSL FLAME. Specifically, we used the FLAME 1+2 approach with automatic outlier deweighting ^96^, which implements Bayesian mixed effects estimation of the group parameter estimates including full Markov Chain Monte Carlo-based estimation for near-threshold voxels ^97^. To identify statistical parametric maps that best represented the average response, all group analyses included age and sex as covariates of no interest (esp. given the developmental sample).

Although our analyses focus primarily on the dorsal attention network (DAN) as the a priori network of interest, we nevertheless conducted whole-brain corrections to the voxelwise GLM statistics to examine the pattern of activity for the signals of interest. Specifically, to correct for familywise error at the whole-brain level, we applied the probabilistic threshold-free cluster enhancement methods ^pTFCE; 98^, thresholding whole-brain maps at *FWE p* < .05 (e.g., Figure 2B). This algorithm provides strict control over familywise error and boosts sensitivity to clusters of activated voxels.

#### Brain-behavior fMRI analyses using regression coefficients from model-based fMRI GLM analyses

To relate individual differences in entropy- and entropy change-related BOLD modulation to behavior on the clock task, we extracted subject-level parameter estimates for these GLM contrasts from each of the 47 DAN parcels defined by the Schaefer cortical parcellation ^see^ ^Table^ ^s1;,34^. These parameter estimates (aka “betas”) served as individual difference measures of sensitivity to signals from SCEPTIC — particularly entropy and entropy change — across regions in the DAN.

We then entered DAN betas for SCEPTIC entropy change as a cross-level moderator of trial-level effects in multilevel models of behavior. Specifically, the dependent variable was trial-wise RT (choice) with behavioral variables as predictors. All models included the trial (inverse-transformed) and previous reward as covariates. Models also included the influence of previous choice (*RT*_t-1_) on current choice (*RT*_t_), or RT autocorrelation. A weaker autocorrelation indicates larger exploratory RT swings, and variables that decrease autocorrelation are considered to increase exploration. Most models included DAN entropy change betas as cross-level moderators of the RT _t-1_ effect as a test of how sensitivity to updates in the number of good options modulated exploration on the task. Likewise, most multilevel models also included the trial-varying location of the best option, *RT*_Vmax_. The two-way interaction of *RT*_Vmax_ and DAN entropy change betas tests whether sensitivity to entropy change enhances or diminishes exploitation of the best option.

Because our behavioral observations had a clustered structure (i.e., trials nested within subjects), we used multilevel regression models, which were estimated using restricted maximum likelihood in the *lme4* package^99^ in *R* 4.2.0^100^. Estimated *p*-values for predictors in the model were computed using Wald chi-square tests and degrees of freedom were based on the Kenward-Roger approximation. For trial-level analyses, subject and run were treated as random and random intercepts were included for these factors. Additionally, as noted in Results, we included random slopes of key terms such as *RT*_Vmax_ and *RT*_t-1_ to ensure the robustness of DAN modulation of exploitation and exploration ^101^.

#### Within-trial mixed-effects survival analyses of behavior with time-varying value estimtaes

To examine the sensitivity of choices to within-trial time-varying value, we performed survival analyses predicting the temporal occurrence of response. These mixed-effects Cox models (R *coxme* package) ^102^ estimated response hazard as a function of model-predicted expected value and their interaction with session-level DAN responses. This survival analysis does not assume that one pre-commits to a given response time, instead modeling the within-trial response hazard function in real, continuous time, accounts for censoring and allows for a completely general baseline hazard function^103^. The survival approach accounts for censoring of later within-trial time points by early responses. Most importantly, it allows for a completely general baseline hazard function that can vary randomly across participants. We thus avoid assumptions about the statistical distribution of response times and account for trial-invariant influences such as urgency, processing speed constraints or opportunity cost. We also modeled only the 1000 – 3500 ms interval, excluding early response times that may be shorter than the deliberation and motor planning period and the end of the interval which one may avoid in order to not miss responding on a trial. We included learned value from the information-compressing model as a time-varying covariate, sampled every 100 ms. To account for between-persons heterogeneity, person-specific intercept was included as a random effects; sensitivity analyses also included the random slope of the predictor of interest (value).

#### Analyses of within-trial peri-feedback BOLD responses using voxelwise deconvolution

Although betas from fMRI GLMs provide a useful window into how decision signals from SCEPTIC relate to behavior at the level of an entire session, the GLM approach makes a number of assumptions: a) that one correctly specifies when in time a signal derived from a computational model modulates neural activity, b) that there is a linear relationship between the model signal and BOLD activity, and c) that a canonical HRF describes the BOLD activity corresponding to a given model-based signal. Furthermore, a conventional model-based fMRI GLM does not allow one to interrogate whether the representation of a given cognitive process varies in time over the course of a trial. For these reasons, we conducted additional analyses that could provide a detailed view of how DAN activity changes following feedback on each trial of the clock task. These analyses also attempted to overcome statistical and conceptual limitations of the GLM and to provide an index of within-trial neural activity that was independent of our computational model. That is, in these analyses, within-trial BOLD activity is the dependent variable and parameters from the SCEPTIC model are predictors.

We first applied a leading hemodynamic deconvolution algorithm to estimate neural activity from BOLD data ^104^. This algorithm has performed better than alternatives in simulated and real fMRI data, and it is reasonably robust to variations in the timing of neural events and the sampling frequency of the scan^105^. We deconvolved the voxelwise BOLD activity for all subjects, averaged the deconvolved time series within each of the 47 DAN parcels (Table s1), and retained these as a regions x time matrix for each run of fMRI data.

Then, to estimate neural activity for each trial in the experiment, we extracted the deconvolved signal surrounding feedback onset (-4 to +4 seconds), censoring timepoints that intersected the previous or next trials. Finally, to ensure that discrete-time models of neural activity could be easily applied, we resampled deconvolved neural activity onto an evenly spaced grid aligned to the feedback onset using linear interpolation. The sampling frequency of the feedback-aligned deconvolved signals was matched to the TR of the fMRI scan (1s for the original sample and 0.6s for the replication sample). Thus, this interpolation was a form of resampling, but did not upsample or downsample the data in the time domain.

For each subject, this yielded a 400 trial x 9 time point (-4—4s for the main sample) x 47 region matrix. We then concatenated these matrices across participants for group analysis. Our primary analyses focused on the four parcels of the DAN visuomotor gradient (MT+, Caudal PPC, Rostral PPC, Premotor), rather than analyzing each region separately. Within each time x parcel combination, we regressed trial-wise neural activity on key decision variables in a multilevel regression framework implemented in *lme4*^99^ in *R*, allowing for crossed random intercepts of subject and side (right/left). Within this framework, the regression coefficients provide an estimate of when and in what region key signals such as entropy change are associated with feedback-related changes in neural activity. Critically, however, given the temporal smoothness of BOLD data, the deconvolved signals remain highly autocorrelated and we are cautious about overinterpreting the temporal precision of these analyses. Moreover, this temporal (and potentially spatial) association results in non-independent statistical tests across the set of space x time models. To adjust for multiple comparisons in non-independent models, we applied the Benjamini–Yekutieli correction across terms of interest in these models to maintain a false discovery rate of .05^106^.

Another advantage of this analytic approach is that alternative models of representation and behavior can be compared in terms of their alignment to neural activity in fMRI. More specifically, each multilevel model across the space x time set of models yields global fit measures such as the Akaike Information Criterion (AIC), which can be used to compare the relative fit of cognitive signals (e.g., entropy change) to event-aligned BOLD data. Here, we used a global model selection approach ^107^ based on the AIC to compare the fit of information-compressing, traditional RL, and working memory accounts of the clock task to activity in the DAN (Figure 3).

### MEG Analyses

#### Multi-level analyses of time-frequency domain MEG data

The goal of these analyses was to estimate how reinforcement modulated oscillatory power at each within-trial timepoint and each frequency. To estimate this effect accurately and robustly across sensors and individuals, we used high-performance parallel computing to fit one multi-level linear model for each point in this time-frequency space, combining data from all trials, individuals, and sensors. Predictors included the SCEPTIC model-derived entropy change signal and behavioral confounds: current and previous response times, reward/omission, trial and, in sensitivity analyses, the KL distance between the last and three preceding response times to account for stochastic choice histories. Subject and sensor were treated as crossed random effects, with sensor-specific random intercepts and random slopes of the behavioral variable of interest and subject-specific random intercepts and, where indicated, random slopes of the variable of interest. Since our contrasts were between trials, the intercept accounted for marginal oscillatory power at a given time-frequency point, and correction for baseline was not necessary. Models were estimated using restricted maximum likelihood in the *lme4* package^99^ in *R* 4.2.0^100^. Estimated *p*-values for predictors in the model were computed using Wald chi-square tests and degrees of freedom were based on the Kenward-Roger approximation. To examine the anatomical distribution of effects, after obtaining estimates for each subject and sensor within this overall model, we projected them into the sensor space (Figure 5C) and source space (Figure 5D) as follows.

Source location was performed using the linearly constrained minimum variance (LCMV) Beamformer procedure^108^. We used Freesurfer’s “fsaverage” template source space and sensor-to-template registration provided by the MNE software^109^. The forward model was calculated using the single-layer boundary element, for a total of 20,484 potential source locations placed with 5 mm spacing on the fsaverage surface. A spatial filter was then constructed using a unit-gain LCMV Beamformer ^108^, with covariances estimated using the 1-second window from the peristimulus interval and 1-second window after the feedback presentation, across all trials and all subjects. We applied the filter to project sensor-level group statistics to the source space. Source estimates were thresholded from 20th to 95th percentiles.

Our analyses of the relationship between subject-level oscillatory responses and behavioral exploration/exploitation employed multi-level survival models identical to those described above (fMRI Analyses, *Within-trial mixed-effects survival analyses of behavior with time-varying value estimates*).

## Author contributions

Conceptualization: AYD, MNH. Data curation: KH, MNH. Formal analysis: AYD, KH, MNH. Funding acquisition: AYD, BL, MNH. Investigation: KH, MNH. Methodology: AYD, KH, MNH. Project administration: AYD, BL, MNH. Resources: AYD, BL, MNH. Software: MNH. Visualization: AYD, KH, MNH. Writing – original draft: AYD, MNH. Writing – review & editing: AYD, BL, KH, MNH.

## Acknowledgments

This work was funded by K01 MH097091, R01 MH067924, and R01MH10095 from the National Institute of Mental Health.

The authors thank Rajpreet Chahal, Mandy Collier, Tanya Shah, Shreya Sheth, Laura Taglioni (data collection), Jiazhou Chen, Bea Langer, and Angela Ianni (data processing and analyses). The authors also thank Carl Olson for helpful comments on an earlier draft of the manuscript.

## Footnotes

1 We note that including entropy, entropy change, and prediction errors simultaneously in GLM analyses does not meaningfully change the pattern of results. This is due to the relatively low level of correlation among these signals.

