## Supplemental Information for "Reinforcement-based option competition in human dorsal stream during exploration/exploitation of a continuous space"

Supplemental figures: 7

Supplemental tables: 4

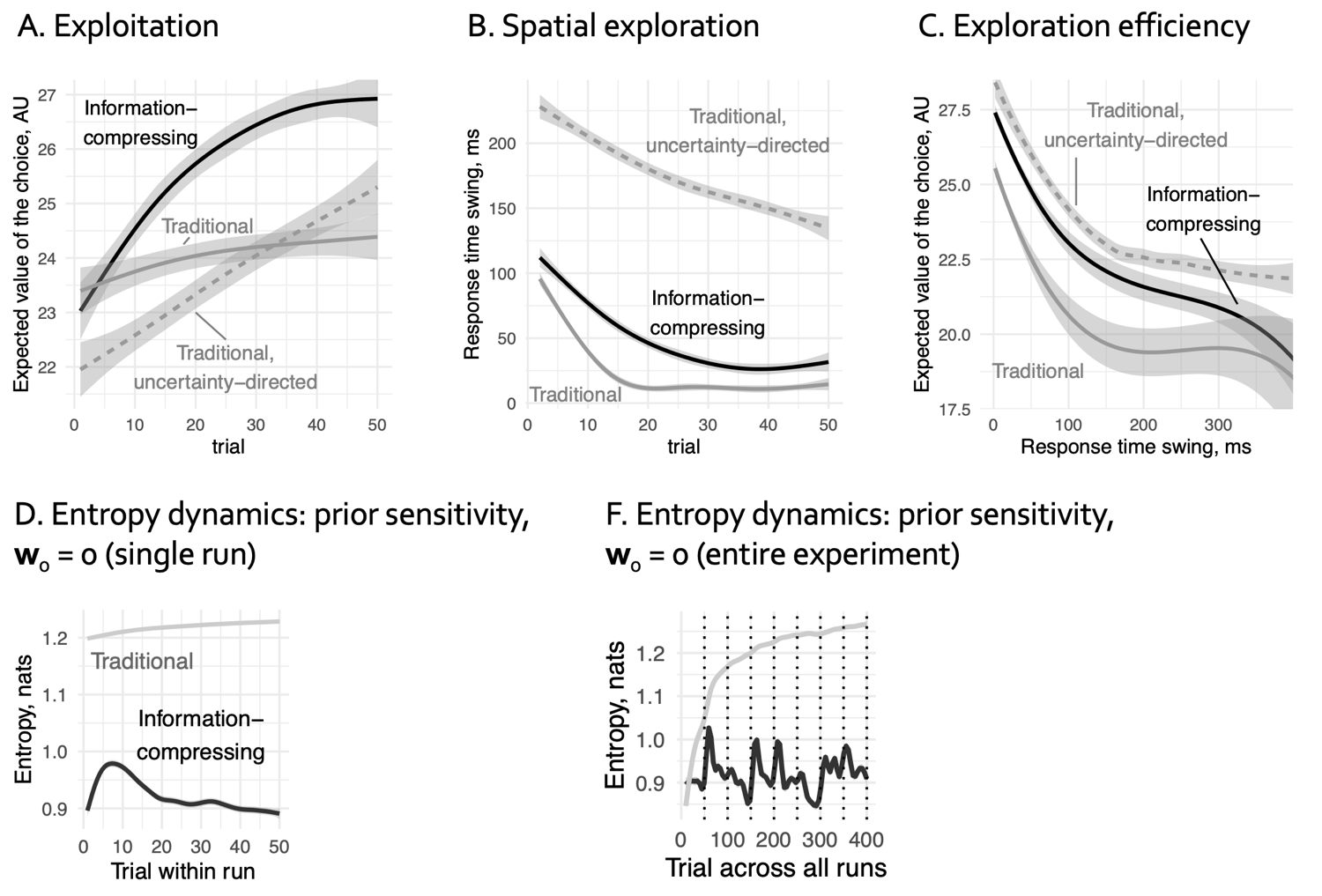

### **Figure s1. Information-compressing RL model: qualitative behavioral predictions and prior sensitivity checks.**

(A) Exploitation: expected reward (value) by trial. Information-compressing (black) and traditional RL (grey, solid) algorithms are compared to more sophisticated but costly uncertainty-directed exploration (grey, dashed). Information-compressing RL model (black) engages in successful exploitation of the continuous space reaching an early, high asymptote. Traditional RL (grey, solid) also asymptotes, but at a lower value. Returns for traditional RL with uncertainty-directed exploration (grey, dashed) continue to improve, but the overall cost of exploration is very high. Data form a computational study described in detail in [Hallquist & Dombrovski, Cognition, 2019](https://psycnet.apa.org/doi/10.1016/j.cognition.2018.11.004) examining model behavior in 40 simulated environments across different parameter sets.

(B) Exploration of the continuous space measured by the absolute distance between consecutive choices (response time swings on the clock task. Traditional RL (grey, solid) decreases its exploration rate early due to long-term value persistence, while information-compressing RL (black) displays more persistent exploration. Uncertainty-directed traditional RL (grey, dashed) displays the highest exploration rate.

(C) Exploration efficiency, illustrating expected reward (ordinate) as a function of the absolute distance between consecutive choices (abscissa). Information-compressing RL (black) explores nearly as efficiently as uncertainty-directed RL (grey, dashed), while traditional RL (grey, solid) explores less efficiently.

(D) Entropy dynamics of the information-compressing vs. traditional RL model across an average run. When prior basis element weights are set to 0 instead of random uniform (cf Figure 1G), the dynamics remain qualitatively unchanged.

(E) The same as D, across the entire experiment (cf Figure 1H).

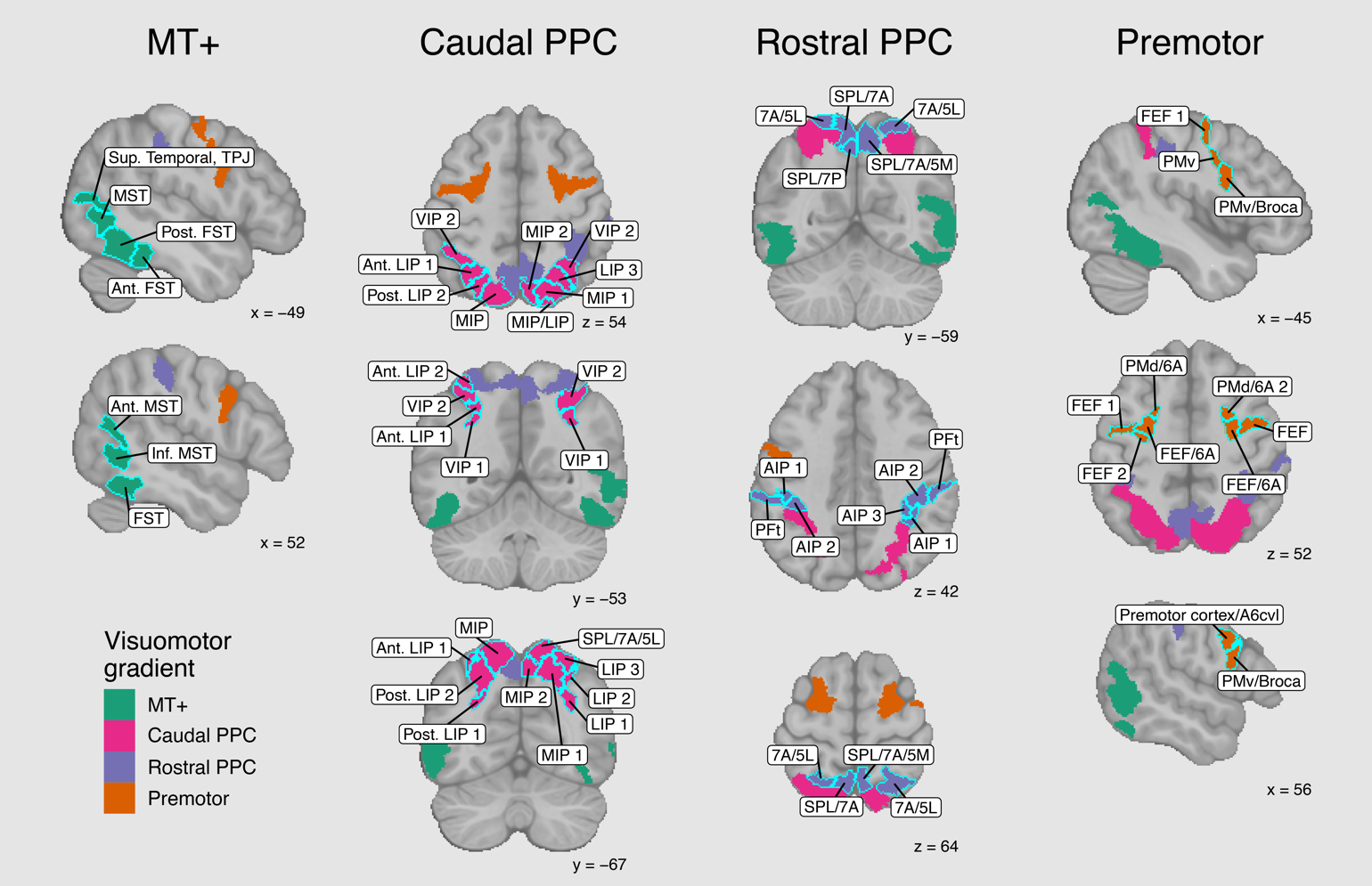

###### **Figure s2. Connectivity-based parcellation of the dorsal stream: individual parcels labeled.**

###### Columns are arranged from late visual to premotor regions. Descriptive labels are provided for comparison to non-human primate studies.

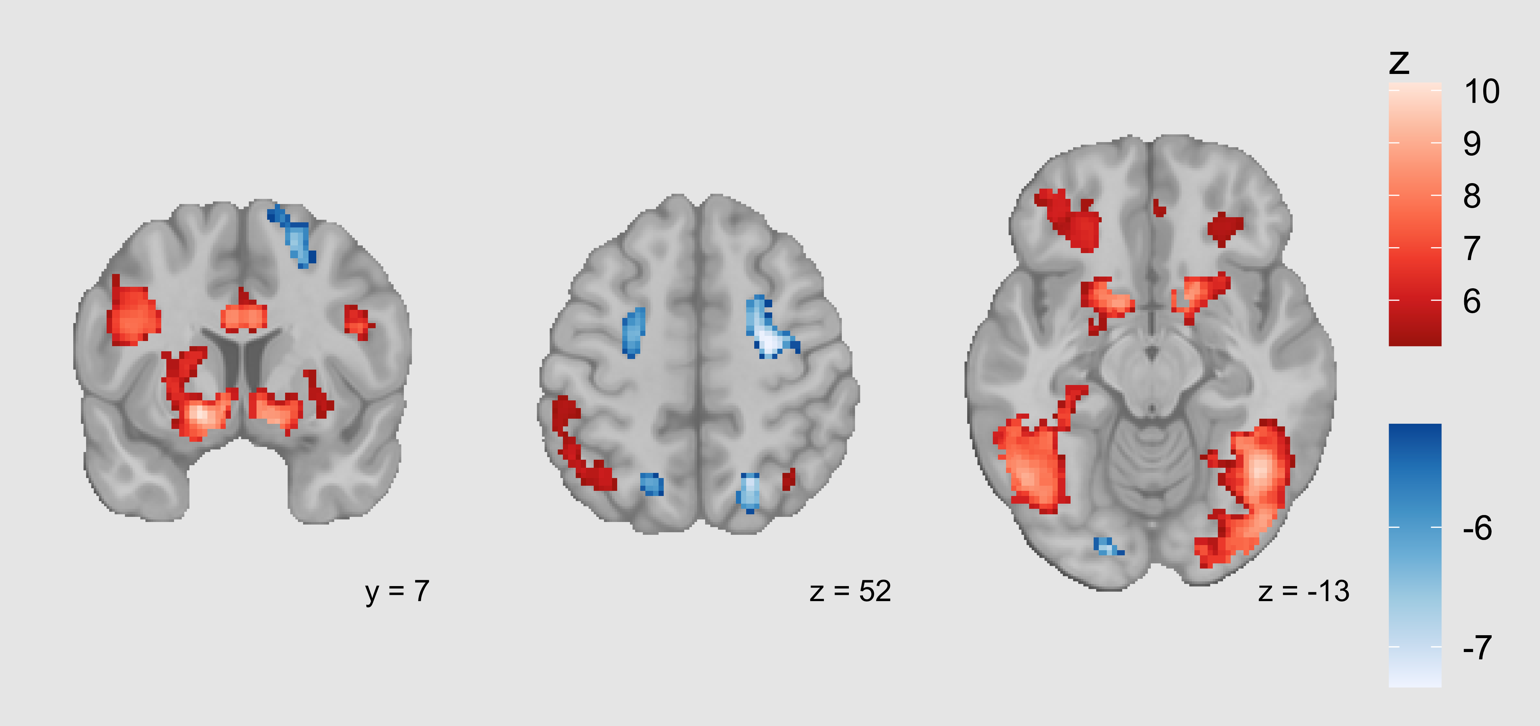

**Figure s3. Responses to reward (red) vs. reward omission (blue), whole-brain analysis.**

As expected, rewards elicit responses in the striatum (left) and orbitofrontal cortex (right), but also in MT+ (right). Reward omissions elicit responses in the frontoparietal nodes of the dorsal attention network (center).

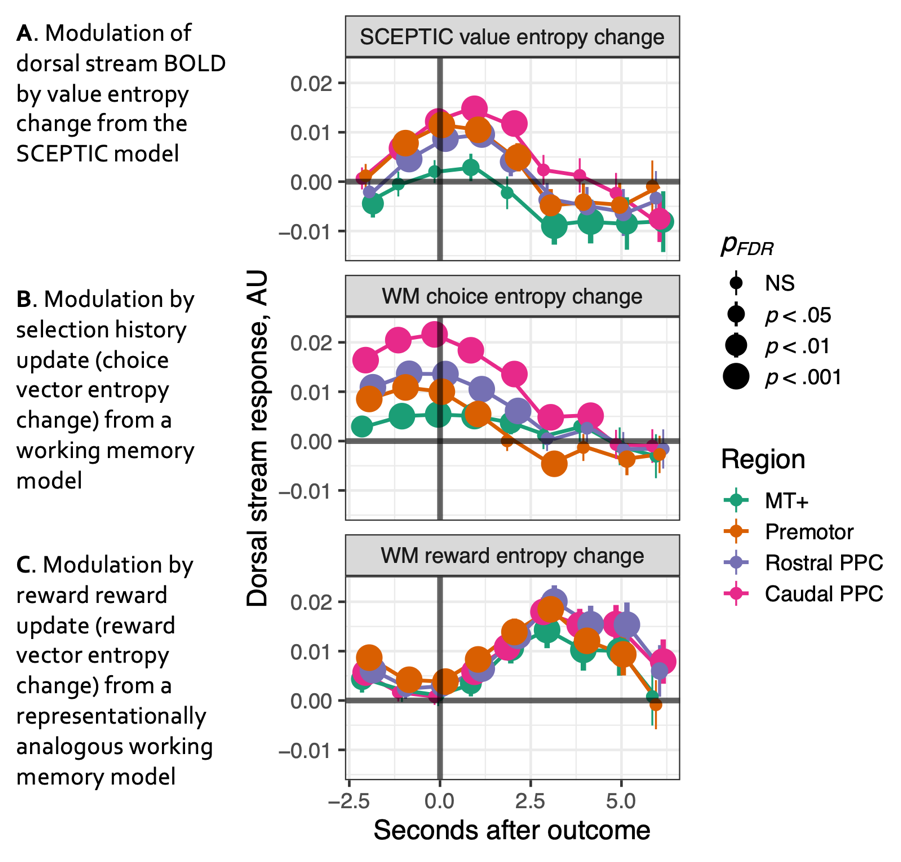

**Figure s4. Information dynamics of the value function, DAN BOLD signal.**

(A) Modulation by SCEPTIC-predicted value entropy change persisted after controlling for the selection history update (B) and reward history update (C) from a representationally analogous working memory model. While BOLD modulation by selection history update occurred peri-response (B), responses to value entropy changed peaked around 1s (A) while responses to reward history update peaked around 3s.

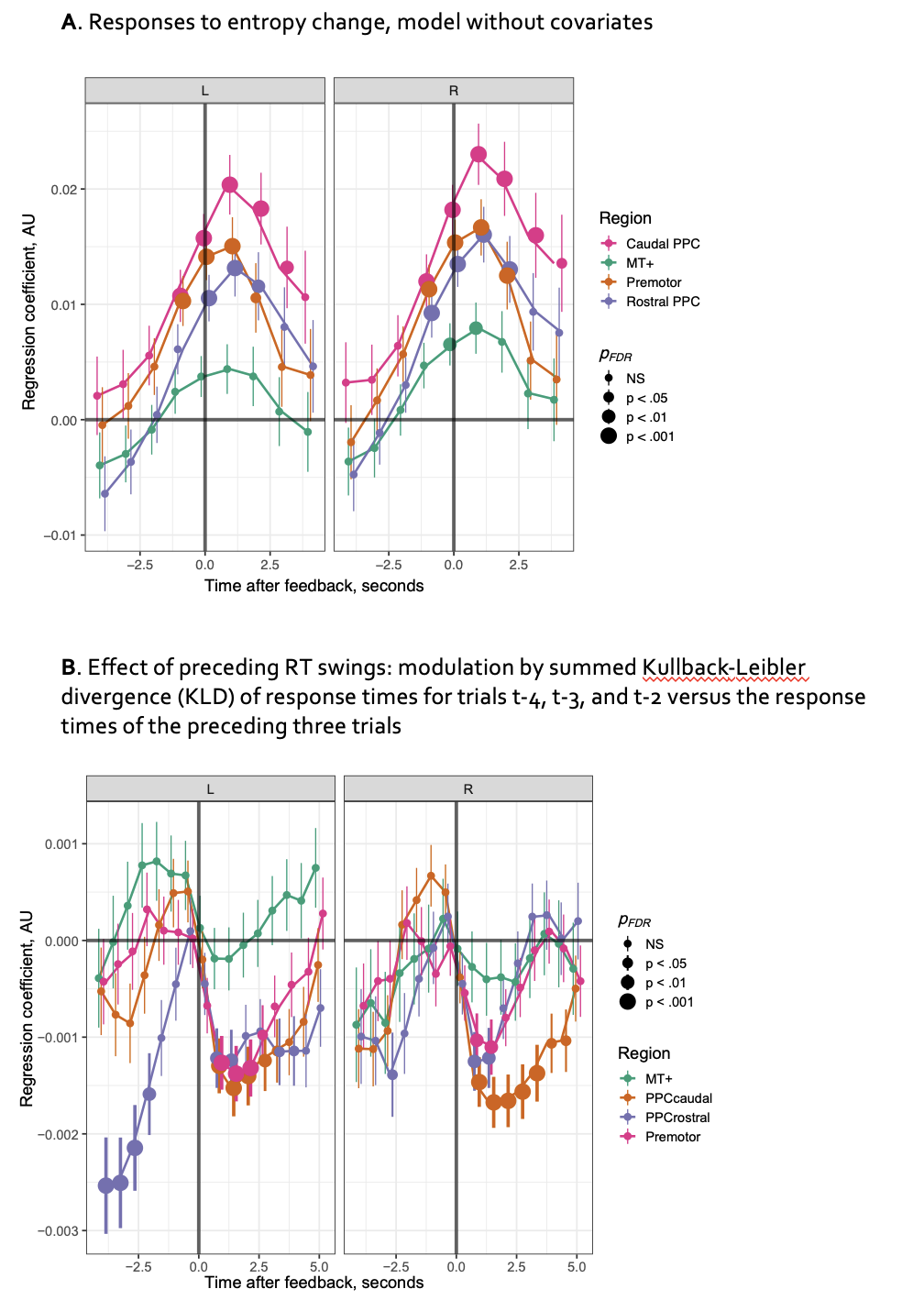

**Figure s5. DAN deconvolved BOLD response to entropy increases: sensitivity analyses**

(A) Model predicting DAN deconvolved BOLD signal with entropy change without controlling for behavioral confounds (current and previous RT, reward/omission, prediction error, V_max,_ entropy).

(B) Selection history may impact value entropy and confound its neural correlates. To rule out this possibility, we computed the summed Kullback-Leibler divergence (KLD) of response times for trials t-4, t-3, and t-2 from the local distribution of response times of the preceding three trials. Higher values of this measure reflect a history of larger RT swings.

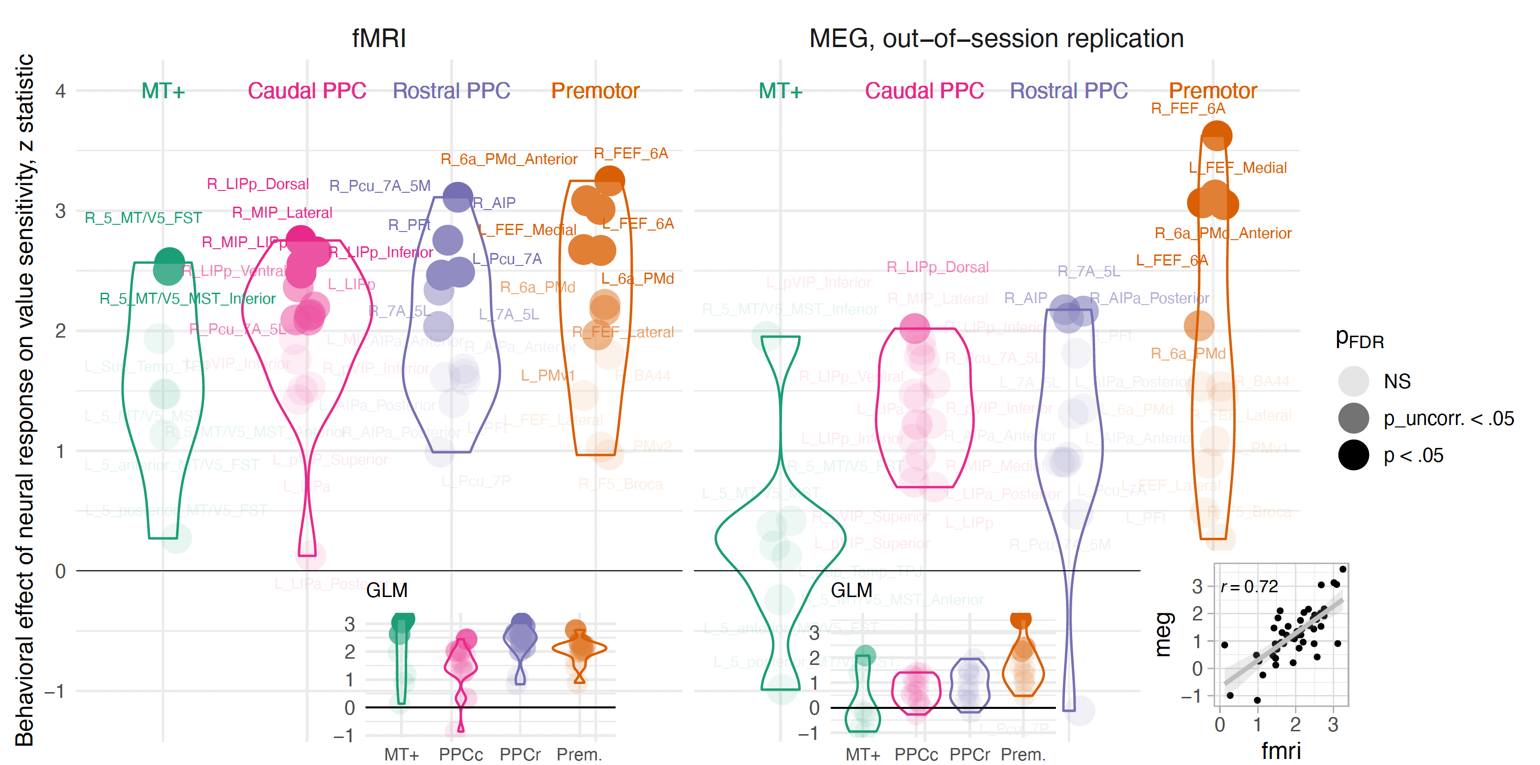

A

B

C

D

E

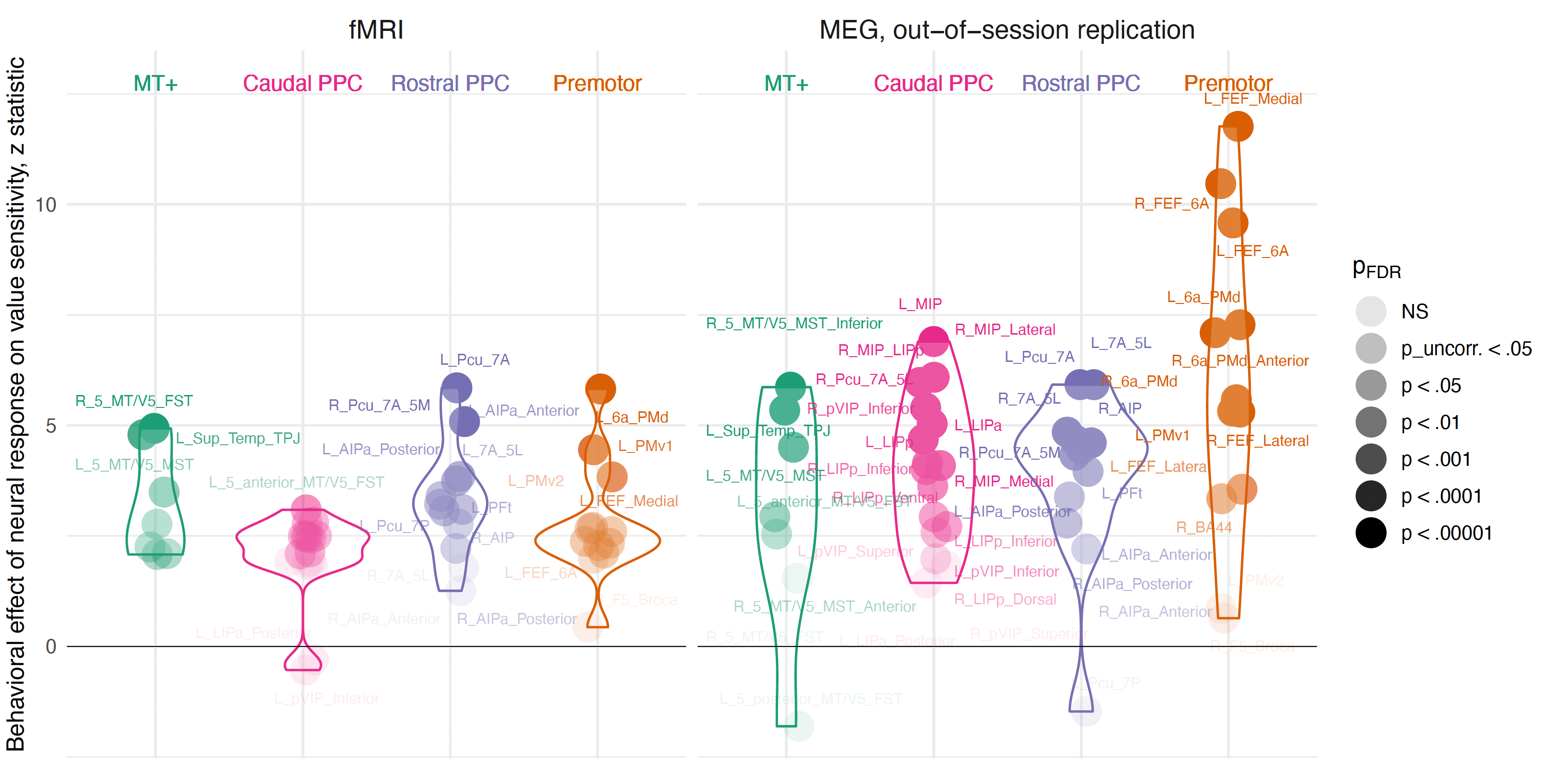

F

G

**Figure s6. BOLD encoding of value information dynamics and behavioral exploitation, sensitivity analyses**

Multi-level survival analyses examining how the individual’s neural response moderates their behavioral sensitivity to within-trial time-varying value. A-D: Models include individual random slopes of value to account for between-persons heterogeneity. F-G: Same model as in Figure 4 but censoring the first 1s and last 0.5s of the interval.

(A) Original fMRI session.

(B) Replication, MEG session. Greater modulation of individual DAN BOLD response by value entropy change predicted more exploitative choices.

(C), (D) Same, GLM analysis.

(E) The anatomical pattern of brain-behavior associations was preserved across the original fMRI and replication sessions. Each dot represents a single DAN parcel as labeled in panels A, B.

(F) Original fMRI session, censoring the first 1s and last 0.5s of the interval.

(F) Replication, MEG session, censoring the first 1s and last 0.5s of the interval.

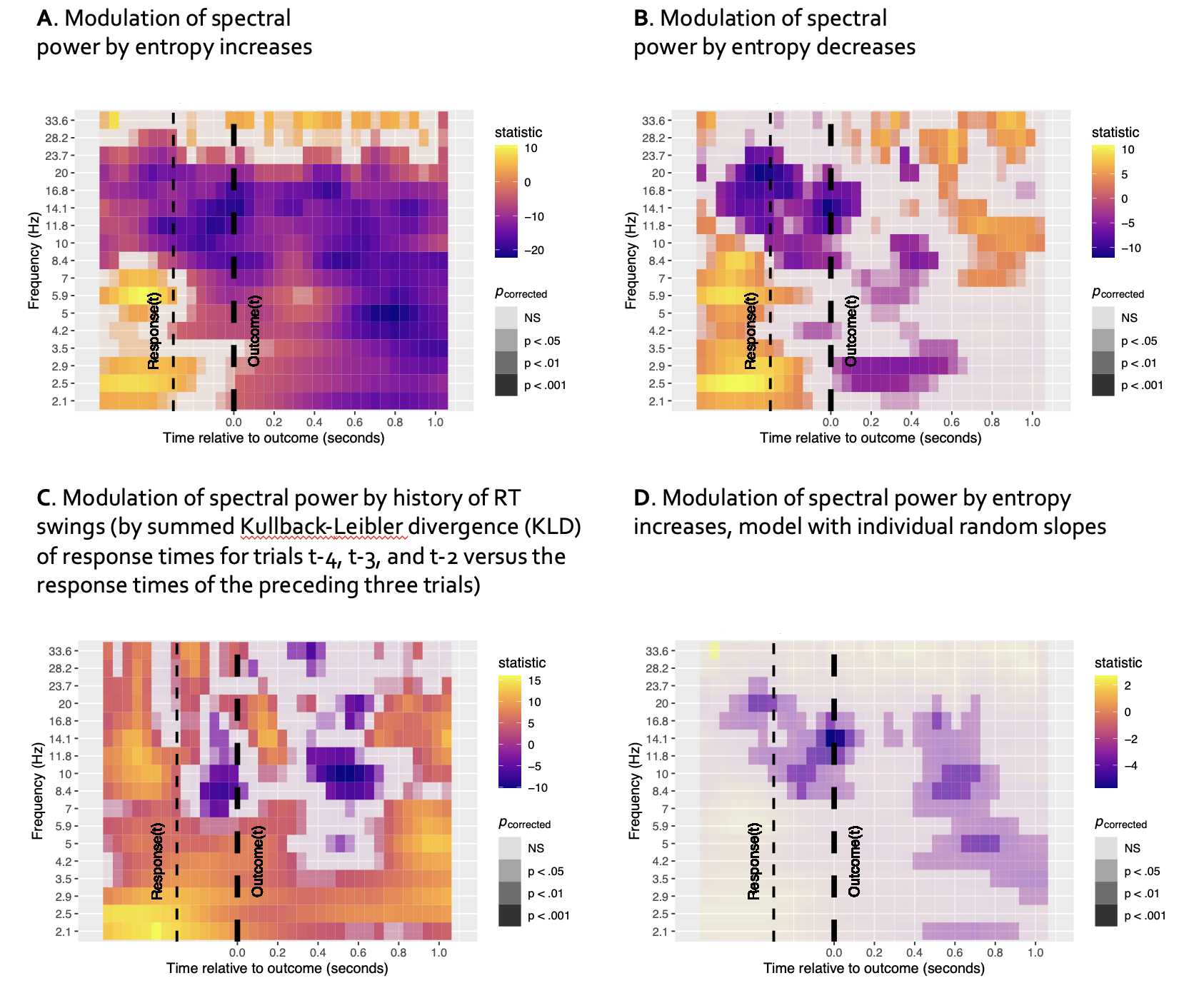
**Figure s7. Suppression of low beta and high alpha oscillatory power in response to entropy increases, sensitivity analyses.**

(A) Oscillatory response to value entropy increases only, with any entropy decrease coded as 0. Cool colors represent de-synchronization to increases reflecting a rising number of potentially valuable options, most prominent between 7-17 Hz (b_1_/a) band at 400-750 ms. The result is nearly identical to the model reported in Figure 5A, which used a full directional entropy change predictor.

(B) Oscillatory response to value entropy decreases only, with any entropy increase coded as 0. The b_1_/a response at 400-750 ms is lacking.

(C) Oscillatory response to RT swings. The The b_1_/a response at 400-750 ms is similar albeit weaker than that elicited by entropy change, suggesting that b_1_/a responses encode both reinforcement and selection history.

(D) Oscillatory response to value entropy change, multi-level model including individual random slopes of entropy change to account for between-subjects heterogeneity.

***Supplementary Table 1.* Schaefer 2018 parcellation of dorsal attention network (DAN) regions**

| **Schaefer node** | **Region label** | **Visuomotor grouping** | **Hemisphere** | **MNI x** | **MNI y** | **MNI z** |
| --- | --- | --- | --- | --- | --- | --- |
| 69 | Anterior FST | MT+ | L | -44 | -42 | -21 |
| 70 | Posterior FST | MT+ | L | -48 | -55 | -15 |
| 71 | MST | MT+ | L | -54 | -62 | -1 |
| 72 | Superior Temporal, TPJ | MT+ | L | -46 | -68 | 14 |
| 73 | Posterior LIP 1 | Caudal PPC | L | -27 | -70 | 29 |
| 74 | PFt | Rostral PPC | L | -53 | -31 | 43 |
| 75 | Posterior LIP 2 | Caudal PPC | L | -24 | -64 | 45 |
| 76 | AIP 1 | Rostral PPC | L | -43 | -30 | 39 |
| 77 | VIP 1 | Caudal PPC | L | -33 | -46 | 39 |
| 78 | Anterior LIP 1 | Caudal PPC | L | -28 | -58 | 48 |
| 79 | AIP 2 | Rostral PPC | L | -36 | -37 | 44 |
| 80 | VIP 2 | Caudal PPC | L | -35 | -50 | 54 |
| 81 | MIP | Caudal PPC | L | -15 | -69 | 56 |
| 82 | Anterior LIP 2 | Caudal PPC | L | -27 | -58 | 60 |
| 83 | SPL/7A | Rostral PPC | L | -7 | -57 | 61 |
| 84 | 7A/5L | Rostral PPC | L | -20 | -54 | 65 |
| 86 | FEF 1 | Premotor | L | -40 | -4 | 51 |
| 87 | FEF/6A | Premotor | L | -26 | -2 | 52 |
| 88 | FEF 2 | Premotor | L | -30 | -9 | 49 |
| 89 | PMd/6A | Premotor | L | -21 | 3 | 62 |
| 90 | PMv/Broca | Premotor | L | -48 | 6 | 25 |
| 91 | PMv | Premotor | L | -49 | 1 | 37 |
| 145 | SPL/7P | Rostral PPC | L | -4 | -62 | 51 |
| 245 | Premotor cortex/A6cvl | Premotor | R | 59 | 6 | 31 |
| 271 | FST | MT+ | R | 48 | -51 | -17 |
| 272 | Inferior MST | MT+ | R | 58 | -56 | -1 |
| 273 | Anterior MST | MT+ | R | 52 | -57 | 12 |
| 275 | LIP 1 | Caudal PPC | R | 31 | -67 | 35 |
| 276 | PFt | Rostral PPC | R | 55 | -25 | 44 |
| 277 | MIP/LIP | Caudal PPC | R | 17 | -76 | 48 |
| 278 | AIP 1 | Rostral PPC | R | 38 | -39 | 38 |
| 279 | AIP 2 | Rostral PPC | R | 43 | -29 | 40 |
| 280 | LIP 2 | Caudal PPC | R | 28 | -61 | 49 |
| 281 | VIP 1 | Caudal PPC | R | 32 | -46 | 41 |
| 282 | MIP 1 | Caudal PPC | R | 20 | -68 | 53 |
| 283 | AIP 3 | Rostral PPC | R | 38 | -38 | 49 |
| 284 | VIP 2 | Caudal PPC | R | 33 | -52 | 53 |
| 285 | MIP 2 | Caudal PPC | R | 7 | -70 | 52 |
| 286 | LIP 3 | Caudal PPC | R | 26 | -58 | 59 |
| 287 | SPL/7A/5M | Rostral PPC | R | 7 | -53 | 57 |
| 288 | SPL/7A/5L | Caudal PPC | R | 13 | -63 | 63 |
| 289 | 7A/5L | Rostral PPC | R | 25 | -51 | 66 |
| 290 | FEF | Premotor | R | 39 | -3 | 51 |
| 291 | FEF/6A | Premotor | R | 27 | -4 | 49 |
| 292 | PMd/6A 1 | Premotor | R | 24 | -4 | 61 |
| 293 | PMv/Broca | Premotor | R | 48 | 8 | 26 |
| 355 | PMd/6A 2 | Premotor | R | 24 | 8 | 57 |

*Note*. These regions were extracted from the 400-node cortical parcellation of Schaefer and colleagues (2018). The nodes included in the DAN differ slightly depending on whether one considers a 7-network or 17-network parcellation. Moreover, we compared DAN nodes in the 200-node variant of the parcellation to ensure robustness of the network definition. The 47 nodes included here are the union of regions contained in the DAN, as defined by the 200- and 400-node solutions, as well as the 7- and 17-network assignments. The Schaefer node numbers reflect the node numbering scheme from the 400-node, 7-network solution. A6cvl = caudal ventrolateral area 6; Brodmann AIP = anterior intraparietal; FEF = frontal eye field; FST = fundus of the superior temporal area; LIP = lateral intraparietal; MIP = medial intraparietal; MT+ = motion-sensitive middle temporal visual area 5; MST = medial superior temporal area; PFt = rostral intraparietal lobule area PFt; PMd = dorsal premotor; PMv = ventral premotor; SPL = superior parietal lobule.

**Table s2.** Clusters of voxels significantly modulated by entropy in whole-brain analyses.

| Label | Network | Hemi-sphere | x | y | z | Voxels | *M z* |
| --- | --- | --- | --- | --- | --- | --- | --- |
| Area 9 Middle | Default | L | -7 | 56 | 8 | 159 | -5.83 |
| Area s32 | Default | R | 5 | 40 | -10 | 136 | -5.64 |
| Area 10r | Default | L | -5 | 54 | -10 | 104 | -5.92 |
| Area a24 | Default | L | -6 | 34 | -9 | 92 | -5.78 |
| Superior 6-8 Transitional Area | Default | R | 23 | 23 | 51 | 60 | 5.90 |
| PreCuneus Visual Area | Default | R | 6 | -48 | 44 | 57 | 6.33 |
| Lateral Area 7P | Dorsal Attention | L | -15 | -69 | 56 | 330 | 8.28 |
| Medial IntraParietal Area | Dorsal Attention | R | 20 | -68 | 53 | 248 | 8.98 |
| PreCuneus Visual Area | Dorsal Attention | R | 7 | -53 | 57 | 191 | 8.15 |
| Anterior IntraParietal Area | Dorsal Attention | R | 38 | -38 | 49 | 167 | 7.45 |
| Area Lateral IntraParietal ventral | Dorsal Attention | R | 26 | -58 | 59 | 151 | 8.26 |
| Area Lateral IntraParietal ventral | Dorsal Attention | R | 33 | -52 | 53 | 128 | 7.47 |
| Medial Area 7P | Dorsal Attention | R | 7 | -70 | 52 | 122 | 9.03 |
| Lateral Area 7A | Dorsal Attention | R | 25 | -51 | 66 | 121 | 7.69 |
| Ventral IntraParietal Complex | Dorsal Attention | L | -27 | -58 | 60 | 108 | 7.06 |
| Medial Area 7A | Dorsal Attention | R | 13 | -63 | 63 | 103 | 9.57 |
| Medial IntraParietal Area | Dorsal Attention | L | -24 | -64 | 45 | 96 | 6.91 |
| Medial Area 7A | Dorsal Attention | L | -7 | -57 | 61 | 96 | 6.98 |
| Area 6 anterior | Dorsal Attention | R | 27 | -4 | 49 | 89 | 7.40 |
| Anterior IntraParietal Area | Dorsal Attention | R | 38 | -39 | 38 | 77 | 7.77 |
| Dorsal Transitional Visual Area | Dorsal Attention | R | 17 | -76 | 48 | 71 | 7.92 |
| Area Lateral IntraParietal ventral | Dorsal Attention | L | -35 | -50 | 54 | 68 | 6.23 |
| Area PGp | Dorsal Attention | R | 43 | -76 | 31 | 68 | 6.54 |
| Area 6 anterior | Dorsal Attention | L | -26 | -2 | 52 | 55 | 5.57 |
| Cerebellum | Frontoparietal Control | L | -31 | -65 | -38 | 345 | 5.93 |
| Area IntraParietal 2 | Frontoparietal Control | R | 51 | -36 | 50 | 134 | 6.65 |
| Medial Area 7P | Frontoparietal Control | L | -4 | -62 | 51 | 133 | 7.14 |
| Inferior 6-8 Transitional Area | Frontoparietal Control | R | 33 | 14 | 54 | 127 | 6.44 |
| Area 6m anterior | Frontoparietal Control | R | 24 | 8 | 57 | 114 | 7.11 |
| Area PFm Complex | Frontoparietal Control | R | 43 | -47 | 47 | 80 | 7.54 |
| Area 7m | Frontoparietal Control | R | 6 | -62 | 44 | 66 | 6.49 |
| Primary Motor Cortex | Somatomotor | L | -34 | -19 | 62 | 95 | -5.39 |
| Primary Sensory Cortex | Somatomotor | L | -32 | -30 | 60 | 53 | -5.37 |
| Cerebellum | Ventral Attention | L | -25 | -56 | -37 | 174 | 5.80 |
| IntraParietal Sulcus Area 1 | Visual | L | -21 | -77 | 43 | 87 | 6.67 |

*Note.* Entropy from the SCEPTIC model was aligned to onset of the decision (clock) phase. Clusters of whole-brain-significant voxels were identified based on a familywise error rate of .05 using the probabilistic threshold-free cluster enhancement (pTFCE) approach. These clusters were then divided into functional regions by calculating the overlap between each data-derived cluster and a whole-brain parcellation adapted from Schaefer and colleagues (2018), detailed in Methods and available here: <https://github.com/UNCDEPENdLab/schaefer_wb_parcellation/blob/main/Schaefer_444_final_2.3mm.nii.gz>. Regions in the parcellation that overlapped with the data-derived clusters by at least 50 voxels are displayed in the table. Spatial coordinates are reported in MNI152 stereotaxic space. Regions are labeled using Glasser and colleagues’ 2016 BALSA parcellation. Voxels were 2.3mm^3^ in size.

**Table s3.** Clusters of voxels significantly modulated by entropy change in whole-brain analyses.

| Label | Network | Hemi-sphere | x | y | z | n Voxels | Mean z |
| --- | --- | --- | --- | --- | --- | --- | --- |
| Superior Frontal Language Area | Default | R | 12 | 18 | 62 | 221 | 8.57 |
| Parieto-Occipital Sulcus Area 1 | Default | R | 12 | -55 | 15 | 284 | 8.75 |
| Parieto-Occipital Sulcus Area 2 | Default | R | 16 | -63 | 28 | 106 | 9.48 |
| IntraParietal Sulcus Area 1 | Dorsal Attention | L | -27 | -70 | 29 | 88 | 9.15 |
| Area PFt | Dorsal Attention | L | -53 | -31 | 43 | 131 | 8.85 |
| Medial IntraParietal Area | Dorsal Attention | L | -24 | -64 | 45 | 194 | 9.35 |
| Anterior IntraParietal Area | Dorsal Attention | L | -43 | -30 | 39 | 65 | 9.32 |
| Area Lateral IntraParietal dorsal | Dorsal Attention | L | -33 | -46 | 39 | 163 | 8.58 |
| Area Lateral IntraParietal ventral | Dorsal Attention | L | -28 | -58 | 48 | 175 | 8.93 |
| Anterior IntraParietal Area | Dorsal Attention | L | -36 | -37 | 44 | 138 | 9.20 |
| Area Lateral IntraParietal ventral | Dorsal Attention | L | -35 | -50 | 54 | 163 | 8.31 |
| Lateral Area 7P | Dorsal Attention | L | -15 | -69 | 56 | 405 | 9.57 |
| Area 6 anterior | Dorsal Attention | L | -26 | -2 | 52 | 99 | 10.30 |
| Area 6 anterior | Dorsal Attention | L | -30 | -9 | 49 | 57 | 9.28 |
| Area 6 anterior | Dorsal Attention | L | -21 | 3 | 62 | 337 | 10.13 |
| Rostral Area 6 | Dorsal Attention | L | -48 | 6 | 25 | 237 | 8.88 |
| Premotor Eye Fields | Dorsal Attention | L | -49 | 1 | 37 | 143 | 8.82 |
| Area IntraParietal 1 | Dorsal Attention | R | 31 | -67 | 35 | 121 | 8.84 |
| Area PFt | Dorsal Attention | R | 55 | -25 | 44 | 163 | 8.51 |
| Dorsal Transitional Visual Area | Dorsal Attention | R | 17 | -76 | 48 | 158 | 9.00 |
| Anterior IntraParietal Area | Dorsal Attention | R | 38 | -39 | 38 | 80 | 8.45 |
| Anterior IntraParietal Area | Dorsal Attention | R | 43 | -29 | 40 | 131 | 8.30 |
| Area Lateral IntraParietal ventral | Dorsal Attention | R | 28 | -61 | 49 | 65 | 8.56 |
| Medial IntraParietal Area | Dorsal Attention | R | 20 | -68 | 53 | 283 | 8.99 |
| Anterior IntraParietal Area | Dorsal Attention | R | 38 | -38 | 49 | 286 | 8.19 |
| Area Lateral IntraParietal ventral | Dorsal Attention | R | 33 | -52 | 53 | 197 | 8.03 |
| Area Lateral IntraParietal ventral | Dorsal Attention | R | 26 | -58 | 59 | 171 | 8.24 |
| PreCuneus Visual Area | Dorsal Attention | R | 7 | -53 | 57 | 242 | 8.34 |
| Frontal Eye Fields | Dorsal Attention | R | 39 | -3 | 51 | 204 | 9.33 |
| Area 6 anterior | Dorsal Attention | R | 27 | -4 | 49 | 92 | 9.77 |
| Area 6 anterior | Dorsal Attention | R | 24 | -4 | 61 | 157 | 9.36 |
| Rostral Area 6 | Dorsal Attention | R | 48 | 8 | 26 | 244 | 8.64 |
| Anterior IntraParietal Area | Frontoparietal Control | L | -45 | -43 | 47 | 131 | 9.00 |
| Anterior Agranular Insula Complex | Frontoparietal Control | L | -34 | 17 | -8 | 121 | 9.48 |
| Area 33 prime | Frontoparietal Control | L | -3 | 4 | 29 | 90 | 8.48 |
| Area IntraParietal 2 | Frontoparietal Control | R | 51 | -36 | 50 | 225 | 7.98 |
| Area PFm Complex | Frontoparietal Control | R | 43 | -47 | 47 | 91 | 8.02 |
| Anterior Ventral Insular Area | Frontoparietal Control | R | 34 | 22 | -6 | 155 | 9.88 |
| Area IFJa | Frontoparietal Control | R | 46 | 18 | 25 | 157 | 9.26 |
| Area 8C | Frontoparietal Control | R | 37 | 11 | 32 | 120 | 8.70 |
| Area 55b | Frontoparietal Control | R | 40 | 7 | 50 | 126 | 9.14 |
| Inferior 6-8 Transitional Area | Frontoparietal Control | R | 33 | 14 | 54 | 286 | 9.01 |
| Area 6m anterior | Frontoparietal Control | R | 24 | 8 | 57 | 151 | 10.19 |
| Area posterior 24 | Frontoparietal Control | R | 8 | 34 | 25 | 267 | 9.58 |
| Area 8BM | Frontoparietal Control | R | 5 | 26 | 47 | 222 | 9.82 |
| Area OP2-3/VS | Somatomotor | L | -36 | -26 | 19 | 135 | 7.81 |
| Area 2 | Somatomotor | L | -53 | -20 | 39 | 129 | 8.44 |
| Primary Sensory Cortex | Somatomotor | L | -47 | -19 | 53 | 159 | 7.95 |
| Area 1 | Somatomotor | L | -46 | -30 | 56 | 250 | 8.80 |
| Area 3a | Somatomotor | L | -37 | -26 | 49 | 45 | 8.28 |
| Dorsal Area 24d | Somatomotor | L | -4 | -9 | 57 | 174 | 8.71 |
| Primary Motor Cortex | Somatomotor | L | -34 | -19 | 62 | 288 | 8.01 |
| Area 2 | Somatomotor | L | -30 | -45 | 60 | 244 | 7.88 |
| Primary Sensory Cortex | Somatomotor | L | -32 | -30 | 60 | 152 | 8.01 |
| Area 6 anterior | Somatomotor | L | -25 | -12 | 61 | 224 | 9.48 |
| Area 6mp | Somatomotor | L | -15 | -13 | 70 | 199 | 7.87 |
| Area PFt | Somatomotor | R | 60 | -15 | 31 | 215 | 8.06 |
| Area 2 | Somatomotor | R | 53 | -17 | 40 | 112 | 8.30 |
| Area 23c | Somatomotor | R | 11 | -16 | 41 | 118 | 7.31 |
| Area 6mp | Somatomotor | R | 27 | -12 | 62 | 159 | 8.36 |
| Anterior Ventral Insular Area | Ventral Attention | L | -32 | 26 | 0 | 138 | 9.73 |
| Frontal Opercular Area 4 | Ventral Attention | L | -32 | 20 | 9 | 159 | 9.08 |
| Frontal Opercular Area 4 | Ventral Attention | L | -43 | 14 | 3 | 128 | 8.44 |
| Anterior 24 prime | Ventral Attention | L | -6 | 21 | 31 | 217 | 8.70 |
| Ventral Area 24d | Ventral Attention | L | -6 | 8 | 47 | 177 | 9.78 |
| Supplementary And Cingulate Eye | Ventral Attention | L | -8 | -5 | 68 | 114 | 8.70 |
| Area TemporoParietoOccipital Junction 1 | Ventral Attention | R | 50 | -41 | 14 | 98 | 7.71 |
| Superior Temporal Visual Area | Ventral Attention | R | 61 | -40 | 22 | 154 | 7.62 |
| Area PF Complex | Ventral Attention | R | 62 | -28 | 39 | 378 | 7.88 |
| Middle Insular Area | Ventral Attention | R | 40 | 9 | -1 | 168 | 8.38 |
| Anterior Ventral Insular Area | Ventral Attention | R | 36 | 23 | 6 | 265 | 9.79 |
| Rostral Area 6 | Ventral Attention | R | 53 | 13 | 12 | 228 | 8.20 |
| Anterior 24 prime | Ventral Attention | R | 7 | 17 | 35 | 291 | 9.14 |
| Area 23c | Ventral Attention | R | 11 | -32 | 42 | 187 | 7.66 |
| Supplementary And Cingulate Eye | Ventral Attention | R | 6 | 9 | 54 | 134 | 9.83 |
| Area 23c | Ventral Attention | R | 10 | -43 | 53 | 116 | 8.52 |
| Supplementary And Cingulate Eye | Ventral Attention | R | 6 | -3 | 65 | 219 | 8.92 |
| Area 6m anterior | Ventral Attention | R | 16 | 4 | 67 | 137 | 9.95 |
| Second Visual Area | Visual | L | -20 | -65 | 7 | 158 | 6.96 |
| Second Visual Area | Visual | R | 16 | -65 | 19 | 223 | 8.62 |

*Note.* Entropy change from the SCEPTIC model was aligned to onset of the feedback, representing change in entropy due to the reinforcement. Clusters of whole-brain-significant voxels were identified based on a familywise error rate of .05 using the probabilistic threshold-free cluster enhancement (pTFCE) approach. These clusters were then divided into functional regions by calculating the overlap between each data-derived cluster and a whole-brain parcellation adapted from Schaefer and colleagues (2018), detailed in Methods and available here: <https://github.com/UNCDEPENdLab/schaefer_wb_parcellation/blob/main/Schaefer_444_final_2.3mm.nii.gz>. Regions in the parcellation that overlapped with the data-derived clusters by at least 75% are displayed in the table. Spatial coordinates are reported in MNI152 stereotaxic space. Regions are labeled using Glasser and colleagues’ 2016 BALSA parcellation. Voxels were 2.3mm^3^ in size.

**Table s4. Effects of reinforcement history on choice beyond the timeframe of the working memory buffer.**

|  | **fMRI session** | | **MEG session** | |
| --- | --- | --- | --- | --- |
| *Predictors* | *Statistic* | *p* | *Statistic* | *p* |
| Intercept | -24.43 | **<0.001** | -32.02 | **<0.001** |
| Trial | -1.17 | 0.242 | -1.60 | 0.109 |
| RT_lag1 | 54.11 | **<0.001** | 64.10 | **<0.001** |
| RT_lag2 | 13.98 | **<0.001** | 18.15 | **<0.001** |
| RT_lag3 | 7.75 | **<0.001** | 6.19 | **<0.001** |
| RT_lag4 | 2.65 | **0.008** | 4.94 | **<0.001** |
| RT_lag5 | 1.73 | 0.084 | 3.92 | **<0.001** |
| Omission_lag1 | 20.47 | **<0.001** | 21.56 | **<0.001** |
| Omission_lag2 | 13.01 | **<0.001** | 12.96 | **<0.001** |
| Omission_lag3 | 6.50 | **<0.001** | 5.55 | **<0.001** |
| Omission_lag4 | 3.09 | **0.002** | 2.67 | **0.008** |
| Omission_lag5 | -0.36 | 0.720 | -1.30 | 0.194 |
| RT_lag1 * Omission_lag1 | -34.49 | **<0.001** | -36.14 | **<0.001** |
| RT_lag2 * Omission_lag2 | -16.00 | **<0.001** | -16.66 | **<0.001** |
| RT_lag3 * Omission_lag3 | -7.38 | **<0.001** | -5.08 | **<0.001** |
| RT_lag4 * Omission_lag4 | -3.79 | **<0.001** | -3.46 | **0.001** |
| RT_lag5 * Omission_lag5 | -0.37 | 0.709 | 0.46 | 0.649 |
| RT_Vmax | 5.42 | **<0.001** | 6.34 | **<0.001** |
| RT_Vmax * Trial | 2.01 | **0.044** | 3.63 | **<0.001** |
| Observations | 26986 | | 32869 | |
| Marginal R^2^ / Conditional R^2^ | 0.228 / 0.363 | | 0.321 / 0.422 | |

*Note.* Statistics from a multi-level model predicting participants’ choices using the working memory buffer and reinforcement history from the information-compressing RL model. RT_lag1-5 represents the selection buffer, and Omission_lag1-5 (vs. reward as reference condition), the reward buffer, and their interactions reflect assignment of rewards to choices. Model-derived RT_Vmax is the location corresponding to the global value maximum. The lack of a RT_lag5 * Omission_lag5 interaction indicates that the buffer does not extend beyond the four preceding trials. A significant effect of RT_Vmax indicates that integrated reinforcement history beyond the timeframe of WM buffer influences choices.
